# *DNAJB1*-*PRKACA* fusion kinase interacts with β-catenin and the liver regenerative response to drive fibrolamellar hepatocellular carcinoma

**DOI:** 10.1101/192104

**Authors:** Edward R. Kastenhuber, Gadi Lalazar, Shauna L. Houlihan, Darjus F. Tschaharganeh, Timour Baslan, Chi-Chao Chen, David Requena, Sha Tian, Benedikt Bosbach, John E. Wilkinson, Sanford M. Simon, Scott W. Lowe

**Author notes:** **Classification**: BIOLOGICAL SCIENCES: Medical Sciences.

## Abstract

A segmental deletion resulting in *DNAJB1-PRKACA* gene fusion is now recognized as the signature genetic event of fibrolamellar hepatocellular carcinoma (FL-HCC), a rare but lethal liver cancer that primarily affects adolescents and young adults. Here, we implement CRISPR/Cas9 genome editing and transposon-mediated somatic gene transfer to demonstrate that expression of both the endogenous fusion protein or a chimeric cDNA leads to the formation of indolent liver tumors in mice that closely resemble human FL-HCC. Notably, overexpression of the wild type PRKACA was unable to fully recapitulate the oncogenic activity of *DNAJB1-PRKACA*, implying that FL-HCC does not simply result from enhanced PRKACA expression. Tumorigenesis was significantly enhanced by genetic activation of β-catenin, an observation supported by evidence of recurrent Wnt pathway mutations in human FL-HCC, as well as treatment with hepatotoxin 3,5-diethoxycarbonyl-1,4-dihydrocollidine (DDC), which causes tissue injury, inflammation and fibrosis. Our study validates the *DNAJB1-PRKACA* fusion kinase as an oncogenic driver and candidate drug target for FL-HCC and establishes a practical model for preclinical studies to identify strategies to treat this disease.

**Significance:** Efforts to understand and treat FL-HCC have been confounded by a lack of models that accurately reflect the genetics and biology of the disease. Here, we demonstrate that the *Dnajb1-Prkaca* gene fusion drives tumorigenesis in mice, and that fusion to DNAJB1 drives FL-HCC initiation more effectively than wild type PRKACA overexpression. The requirement of the PRKACA kinase domain in tumor initiation establishes the potential utility of kinase inhibitors targeting the fusion. By identifying genetic and environmental factors that can enhance the consistency and aggressiveness of disease progression, we reveal biological characteristics of the disease and advance a robust platform for future pre-clinical studies.

## Introduction

Fibrolamellar hepatocellular carcinoma (FL-HCC) ubiquitously harbors a ~400kb deletion on chromosome 19 that produces an in-frame fusion of the DnaJ heat shock protein family member B1 (*DNAJB1)* and protein kinase cAMP-activated catalytic subunit alpha (*PRKACA)* (**Fig. S1A**) (1, 2). *DNAJB1* encodes a subunit of the heat shock factor 40 (HSP40) complex, which activates the ATPase of HSP70 and serves as a molecular chaperone that can be induced by an array of environmental stresses (3). *PRKACA* encodes a catalytic subunit of protein kinase A (PKA), which resides in the cytoplasm in an inactive tetrameric complex with PKA C-β and two regulatory subunits of the PKA holoenzyme (4). Activation of GPCRs leads to cAMP-dependent activation of the PKA catalytic subunits and subsequent phosphorylation of a panoply of cellular substrates (4). The crystal structure of the DNAJB1-PRKACA fusion protein shows that the catalytic site, regulatory subunit binding and anchoring protein binding remain similar to the wild type PRKACA (5).

Beyond the presence of *DNAJB1-PRKACA* fusions, FL-HCC tumorigenesis is poorly understood. Few, if any, other significantly recurrent mutated genes have been described (6, 7), and while broad copy number alterations have been observed, they do not specifically implicate known oncogenes or tumor suppressors (7). Unlike liver cancer in older adults, FL-HCC is not associated with any known etiological risk factors such as alcoholism, chronic hepatitis infection, or liver flukes (8).

Currently, FL-HCC is diagnosed on the basis of histological features such as large cells with granular eosinophilic cytoplasm, vesiculated nuclei, and large nucleoli. Ultrastructural studies observe a hyperaccumulation of mitochondria and abundant endoplasmic reticulum (9). While early onset and lack of chronic liver disease are suggestive of FL-HCC, classic HCC can also occur in young patients and misdiagnosis is common (10). Given the specificity of *DNAJB1-PRKACA* fusion for FL-HCC, its detection will likely be decisive for correct diagnosis (2).

Surgical resection is currently the primary treatment for FL-HCC patients. Although often described as a relatively indolent disease, a high rate of recurrence represents a major clinical challenge (11) and the 5-year survival rate is 34% (12). There is no evidence of survival benefit from adjuvant chemotherapy or any systemic treatment applicable to classic HCC (13). While the unique demographics and genetics of FL-HCC suggest that these patients should be treated differently than those with HCC, there have been few clinical trials that have been tailored to this patient population (www.clinicaltrials.gov). The development of FL-HCC-specific therapies has been further hindered by the lack of genetically and biologically accurate model systems.

Mouse models have been a powerful tool to evaluate the oncogenic potential of candidate drivers, to study the biology of tumorigenesis, and as preclinical systems to test novel therapeutics (14). In this study, we employed hydrodynamic transfection combined with either CRISPR/Cas9-mediated editing of the endogenous deletion or transposon-mediated transgenesis of fusion cDNA and variants, allowing us to introduce genetic lesions in a subset of hepatocytes without the time and expense of producing germline genetic strains (15, 16). Using this approach, we demonstrate that the *DNAJB1-PRKACA* fusion is a *bona-fide* oncogene and identify genetic and environmental factors that cooperate with the fusion event to drive aggressive disease. Our results further show that the PRKACA kinase domain is required for these effects, providing rationale for targeting kinase activity pharmacologically. We anticipate that the models presented herein will serve as a powerful platform for future biological and pre-clinical studies.

## Results

### CRISPR-mediated deletion results in fusion oncogene and drives tumorigenesis in vivo

The oncogenic potential of the endogenous *DNAJB1-PRKACA* fusion *in vivo* was assessed. Co-expression of Cas9 with multiple single guide RNAs (sgRNAs) can be used to model chromosome translocations, inversions, and deletions by generating DNA double strand breaks (DSBs) at the breakpoints of chromosome rearrangements, which are subsequently joined by non-homologous end-joining (NHEJ) (17-21). While such events are rare, an oncogenic rearrangement is expected to be positively selected *in vivo*. We determined whether the FL-HCC-associated rearrangement could be generated in hepatocytes of young adult mice via hydrodynamic tail vein injection using tandem sgRNAs corresponding to the breakpoints of the disease-associated deletion in the first introns of *Dnajb1* and *Prkaca* (**Fig. 1A**). Importantly, the deleted region on human chromosome 19 in FL-HCC is syntenic to a corresponding region on mouse chromosome 8. In fact, all protein-coding genes present in the human region have orthologs present in the mouse region, and are arranged in the same order (**Fig. 1A**).

**Fig. 1.**
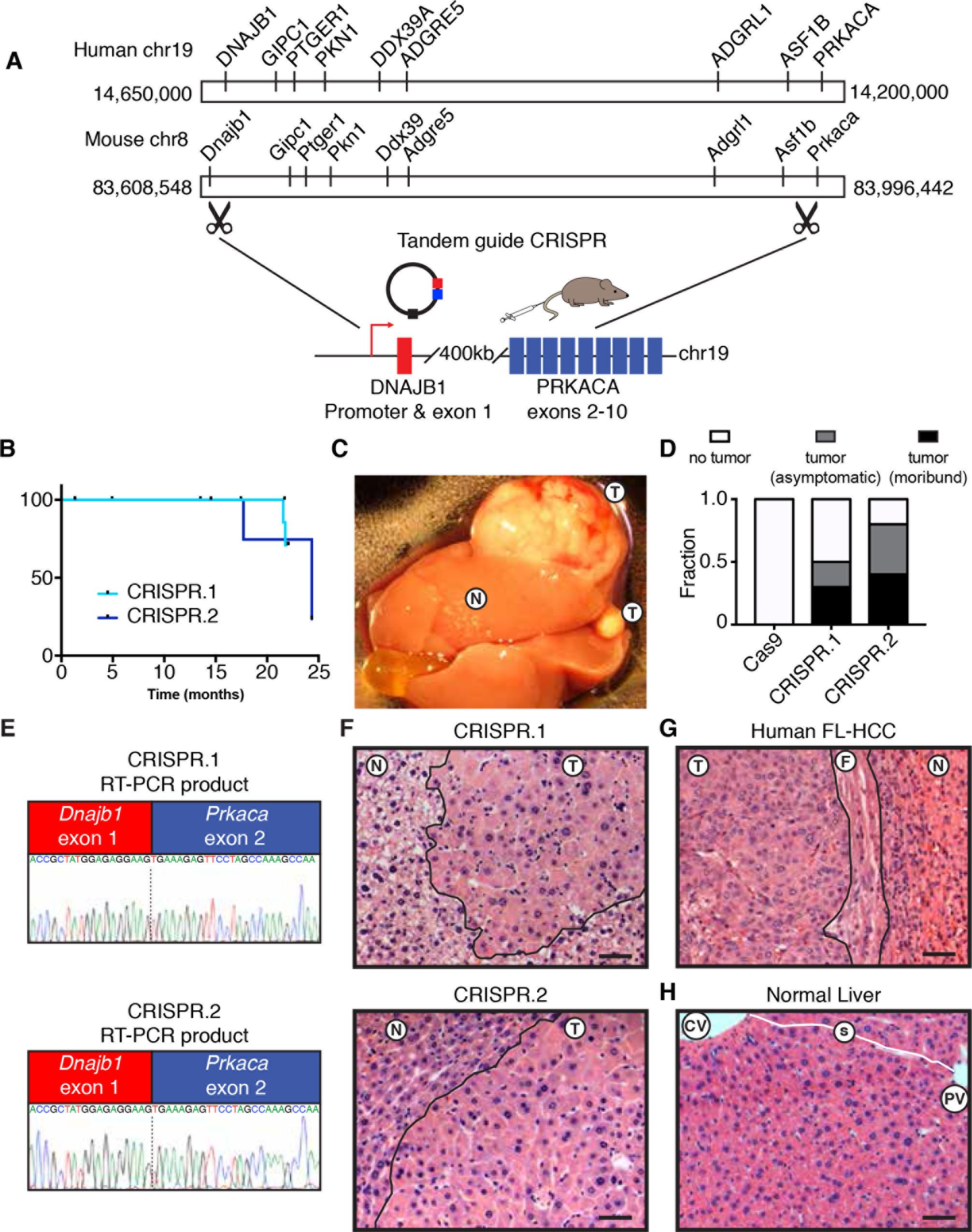
CRISPR-mediated deletion results in fusion oncogene and drives tumorigenesis in vivo. (**A**) Configuration of human chromosome 19, including *DNAJB1* and *PRKACA*, configuration of mouse chromosome 8, including *Dnajb1* and *Prkaca*, and schematic of endogenous 400kb deletion targeted by hydrodynamic injection of vector containing tandem guide sgRNAs to introns of *Dnajb1* and *Prkaca* and Cas9. (**B**) Overall survival of mice injected with CRISPR.1 (n=10) or CRISPR.2 (n=9). (**C**) Macroscopic view of a tumor-bearing liver (**D**) Fraction of mice harvested with no detectable tumor, asymptomatic mice with histologically detectable disease (asymptomatic), or moribund mice with tumors (moribund) for each indicated genotype. (**E**) Sanger sequencing of chimeric transcript amplified from tumors generated by CRISPR.1 (top) and CRISPR.2 (bottom). (**F**) H&E staining of tumor generated by CRISPR.1 (top) and CRISPR.2 (bottom). (**G**) Human case of FL-HCC (T, tumor; F, fibrosis). (**H**) Normal mouse liver, where sinusoids trace from central veins to portal triads (CV, central vein; PV, portal vein) with intact sinusoids (white line, s). All scale bars are 50 αm.

To test the feasibility of this approach, different sgRNAs capable of targeting the first intron of *Dnajb1* and *Prkaca* were co-expressed with Cas9 in NIH3T3 cells or adult livers using lentiviral transduction or hydrodynamic injection, respectively (**Fig. S1A**), and confirmed to produce a fusion event using PCR (**Fig. S1B, C**). Next, two sgRNA pairs, targeting different sequences within the same introns (herein CRISPR.1 and CRISPR.2), were introduced into the livers of adult mice and the animals were monitored over time. A subset of animals transduced with both sgRNA combinations became moribund with liver tumors 16-24 months post injection (**Fig. 1B**). Tumor-bearing mice typically harbored disease involving multiple lobes, presumably from independent initiating events and ranged from diffuse to macroscopically visible (**Fig. 1C**). In samples evaluated histologically, 2/9 mice injected with CRISPR.1, and 3/7 mice injected with CRISPR.2, died as a result of tumor burden. Additionally, non-moribund animals that were sacrificed harbored histological evidence of disease (annotated as “asymptomatic”) in 2/9 CRISPR.1 mice and 2/7 of CRISPR.2 mice (**Fig. 1D**). For both guide pairs, RT-PCR and Sanger sequencing with fusion-specific primers confirmed expression of the intended *Dnajb1-Prkaca* fusion oncogene in these lesions (**Fig. 1E**).

Histologically, the CRISPR-induced mouse tumor cells (**Fig. 1F**) were strikingly similar to human FL-HCC (**Fig. 1G**). Like human FL-HCC, the mouse liver tumors were composed of large, pleiomorphic polygonal cells with abundant eosinophilic cytoplasm, large vesicular nuclei and prominent nucleoli. Furthermore, the lobular structure of the tumors was clearly disrupted (**Fig. 1H**) and distinctive cytoplasmic inclusions were observed (see below). However, unlike the human disease, the mouse tumors were not surrounded by detectable fibrosis. Supporting the robustness of these results, tumors with similar latency and histology were recapitulated with an independent Cas9-expressing vector (**Fig. S1D**). Thus, induction of an endogenous *Dnajb1-Prkaca* fusion through intrachromosomal deletion drives tumors with features of FL-HCC in mice. Independent work from others recently demonstrated an FL-HCC phenotype of lesions in asymptomatic mice using similar methods (22).

### *DNAJB1-PRKACA* fusion drives liver tumorigenesis

The segmental deletion that results in the *DNAJB1-PRKACA* gene fusion entails heterozygous loss of 7 other coding genes, with unknown functional contribution. In other contexts, such deletions can contribute to tumorigenesis directly through attenuating the function of haploinsufficient tumor suppressors (23). To determine whether the *DNAJB1-PRKACA* fusion is sufficient to drive tumorigenesis (uncoupled from the typical genomic deletion), and whether simply the overexpression of the wild-type PRKACA gene could recapitulate this effect, we used hydrodynamic injection to deliver a transposon expressing the human *DNAJB1-PRKACA* fusion cDNA or a full length wild-type *PRKACA* cDNA. Co-transfection of transiently expressed sleeping beauty transposase (“SBase”) with a transposon construct allows for stable integration and constitutive overexpression of the cDNA that mimics the high levels in human tumors (1) (**Fig. 2A**).

**Fig. 2.**
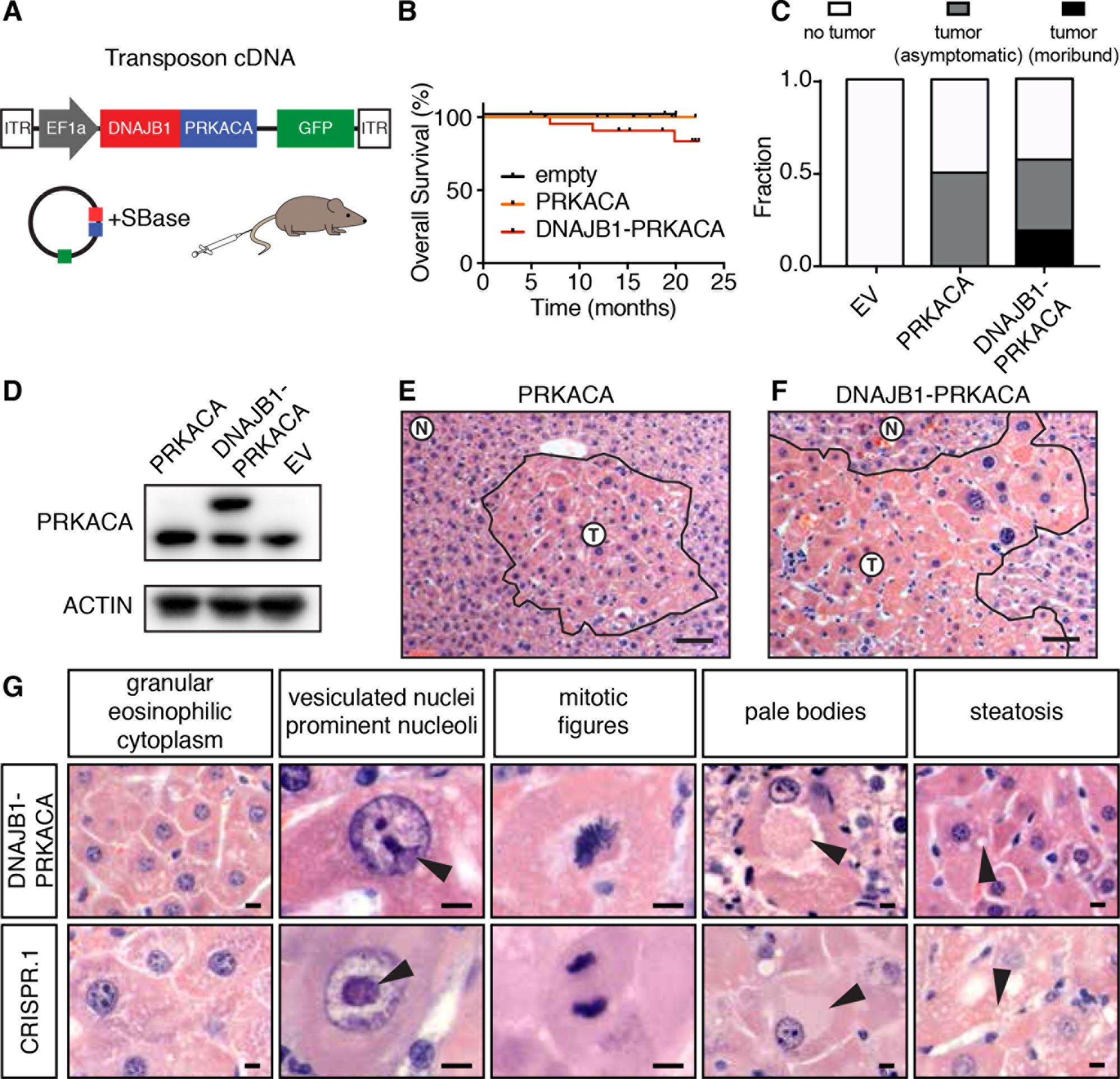
*DNAJB1-PRKACA* fusion drives liver tumorigenesis. (**A**) Schematic of Sleeping Beauty transposon strategy to deliver *DNAJB1-PRKACA* fusion cDNA to young adult livers (ITR: inverted terminal repeats). (**B**) Overall survival of mice injected with cDNA encoding *DNAJB1-PRKACA* (n=23), wild-type *PRKACA* (n=12), or empty vector (n=4). (**C**) Fraction of mice harvested with no detectable tumor, asymptomatic mice with histologically detectable disease, or moribund mice with tumors expressing empty vector (EV), wild type *PRKACA*, or *DNAJB1-PRKACA* fusion. (**D**) Western blot of liver progenitor cells 4 days after transduction with indicated constructs *in vitro* (**E**) Cluster of atypical hepatocytes in liver injected with wild type pT3-PRKACA (**F**) Tumor generated by pT3-DNAJB1-PRKACA (T, tumor; N, adjacent normal; scale bar=50um). (**G**) Higher magnification image highlighting common murine FL-HCC features: large granular, eosinophilic cytoplasm; vesiculated nuclei with prominent nucleoli; mitotic figures; pale bodies; steatosis. Scale bars=10 αm.

Expression and protein stability are similar between the DNAJB1-PRKACA fusion protein and full-length WT PRKACA, and overexpression of wild type PRKACA produced some changes to hepatocyte histology, but it did not trigger the formation of lethal tumors (**Fig. 2B-E**). However, expression of the *DNAJB1*-*PRKACA* cDNA produced tumors with similar kinetics, penetrance, and morphology as CRISPR-driven murine FL-HCC (**Fig. 2B,C,F**). Again, a spectrum of histological findings supported the similarity between these complementary methods and human FL-HCC (**Fig. 1,2**), including the presence of large tumor cells with granular, eosinophilic cytoplasm and prominent nucleoli (**Fig. 2F**). Mitotic figures, steatosis, and pale bodies (8) were also observed (**Fig. 2G**). These results imply that the DNAJB1 portion of the fusion protein contributes to disease beyond facilitating overexpression of PRKACA and that the chromosome 19 deletion event is not required for oncogenesis.

### Murine tumors display structural and molecular features of human FL-HCC

To validate that the murine model recapitulates other aspects of the human disease, we further characterized the phenotype of murine tumors. Human FL-HCC has consistently recognizable ultrastructural features that were also observed in murine FL-HCC tumors by electron microscopy (EM). Like human FL-HCC, murine tumor cells were typically larger than adjacent normal hepatocytes and contained clumped heterochromatin (**Fig. 3A-B**,**S2A**), occasional pale bodies (**Fig. 3B**), and prominent nucleoli (**Fig. S2B,C**). Most notably, tumor cells exhibited a marked increase in mitochondria with atypical appearance (yellow arrows, **Fig. 3B,C,D**,**S2E-H**) (8, 9, 24). Numerous megamitochrondria were observed. The mitochondria were round to oval and homogeneous without obvious cristae and were surrounded by abundant rough endoplasmic reticulum (red arrows, **Fig. 3C,D**,**S2**). Throughout the tumor, cells had moderate to severe perinuclear and cytoplasmic aggregates of lipofuscin pigment (**Fig. 3B,S2I**). This finding, along with the mitochondrial phenotype, could be consistent with a state of oxidative stress (25, 26).

**Fig. 3.**
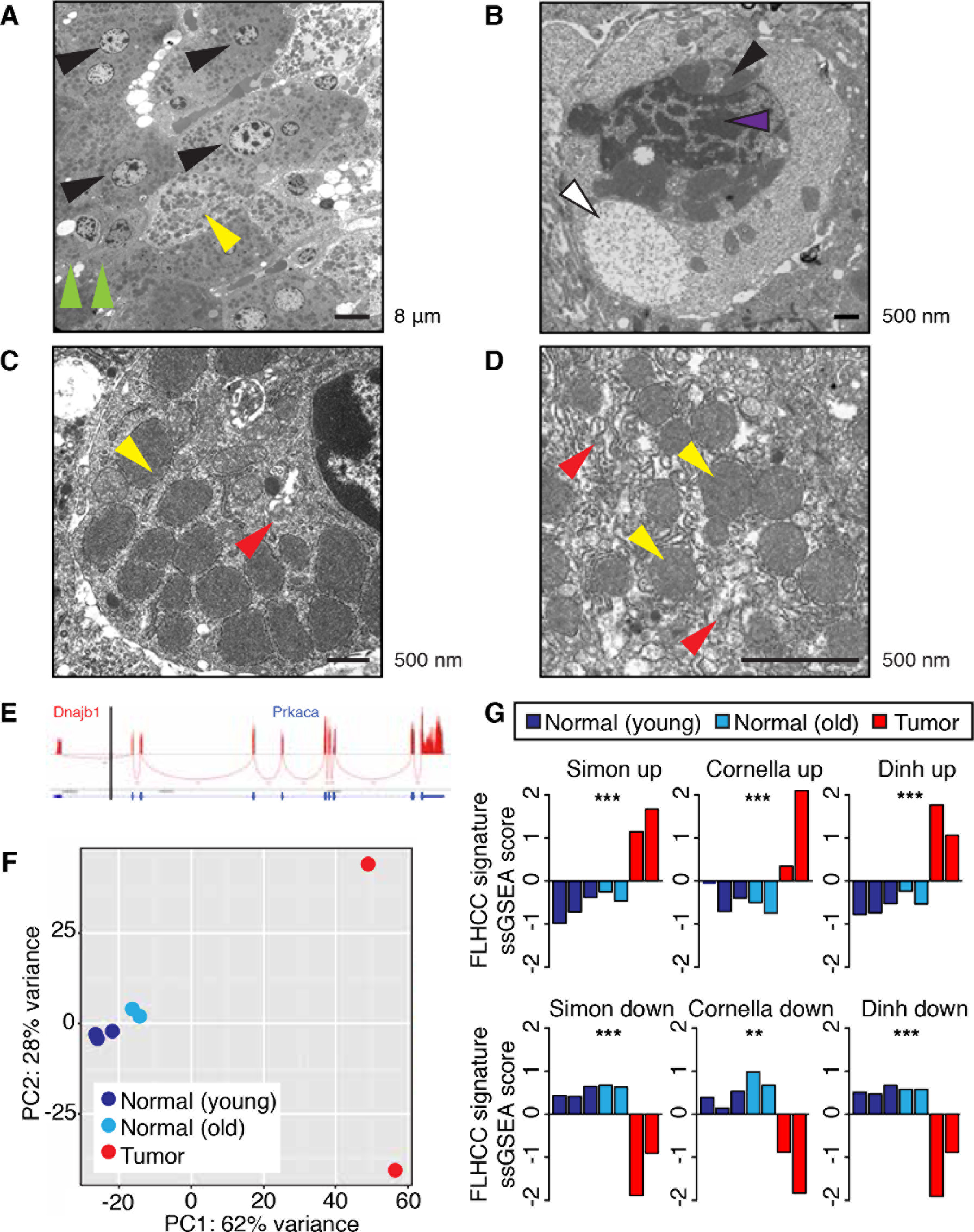
Murine tumors display structural and molecular features of human FL-HCC. Ultrastructural analysis of a transposon-induced tumor by electron microscopy (**A**) Tumor cells (black arrows) of varying larger size than compressed adjacent hepatocytes at tumor margin (green arrows). (**B**) Pale body (white), perinuclear accumulation of lipofuscin (black) and heterochromatin (purple). (**C**) Tumor cell with abundant mitochondria (yellow), Rough endoplasmic reticulum (RER, red). (**D**) Abnormal mitochondria (yellow) with indistinct cristae surrounded in RER (red). (**E**) Sashimi plot indicating RNA-seq reads that cross the *DNAJB1-PRKACA* junction. (**F**) Principal component analysis of young control livers (dark blue), aged control livers (light blue), and tumors derived from CRISPR.1 (red). (**G**) Z scored single sample gene set enrichment analysis (ssGSEA) for normal (black) and tumor (red) samples. **P<0.01 ***P<0.001.

Human FL-HCC is known to often express markers of multiple lineages, including hepatic, billiary, and neuroendocrine (27, 28). The murine tumors were positive for hepatocyte markers HNF4A and HNF1A, with some cells showing reduced expression, consistent with reduced hepatocyte lineage commitment (**Fig. S3A,B**). However, the murine tumors were negative for other proteins that are often expressed in FL-HCC, including biliary markers CK7 and CK19 as well as CD68 (**Fig. S3C-F**), perhaps reflecting the fact that mature hepatocytes are targeted by the hydrodynamic transfection technique and thus are necessarily the cell of origin of the murine FL-HCC model, whereas the cell of origin in human FL-HCC is unknown.

To further validate the mouse model, gene expression analysis by RNA-Seq was performed on murine tumors and control liver tissue (**Table S1**) and compared to human FL-HCC. Sequencing reads that cross the junction were observed, confirming expression of the fusion (**Fig. 3G**). Principal component analysis demonstrated that the vast majority of the variance between samples described the differences between tumor and normal samples (**Fig. 3H**). A focused analysis to investigate the similarity between mouse and human tumors was evaluated in two ways. First, single sample gene set enrichment analysis (ssGSEA) (29), using FL-HCC expression signatures from three independent published studies (7, 10, 30), was used to confirm that differentially expressed genes in human tumors were, in aggregate, significantly enriched in our murine tumors (**Fig. 3I**). Second, a supervised analysis of curated functional gene sets previously reported as enriched in FL-HCC (30) was consistent with murine tumor expression data (**Fig. S4A**). These results provide a global analysis that classifies the murine model as FL-HCC. Overall, the murine tumors arising in the presence of the *DNAJB1-PRKACA* fusion show most, but not all, features of the human disease.

### Molecular profiling of murine FL-HCC reveals processes linked to tumorigenesis

Unbiased analysis of expression data also suggested yet other biological processes that may be relevant to FL-HCC pathogenesis. A total of 5710 genes were significantly differentially expressed between tumor and normal tissue (**Fig. 4A**, **Table S2**) and further global analysis by GSEA was performed (**Fig. 4B**, **Table S3**). The gene expression data, in agreement with human signatures, showed evidence of increased proliferation and mitogenic signaling (**Fig. 4A-C**). For example, cell cycle and DNA biosynthesis gene sets, and specifically *Cdk1*, *Gins2*, and *Cenpa,* were highly upregulated in the experimental tumors (**Fig. 4D,E**). Accordingly, tumors displayed an elevated Ki67 index (~9%) compared to adjacent normal tissue (~1%) (**Fig. 4C**). Activation of the PI3K pathway was indicated by downregulation of *Deptor* (a negative regulator of mTORC1 also decreased in human tumors) and upregulation of the RTK ligands *Egf*, *Nrg2*, and *Ereg* in both mouse and human tumors (30) (**Fig. 4A**,**S4E**). This observation was validated by immunofluorescence showing high levels of phospho-S6rp, a marker of mTOR activity that is highly expressed in most FL-HCCs but rarely in classic HCC (7, 31) (**Fig. 4C**).

**Fig. 4.**
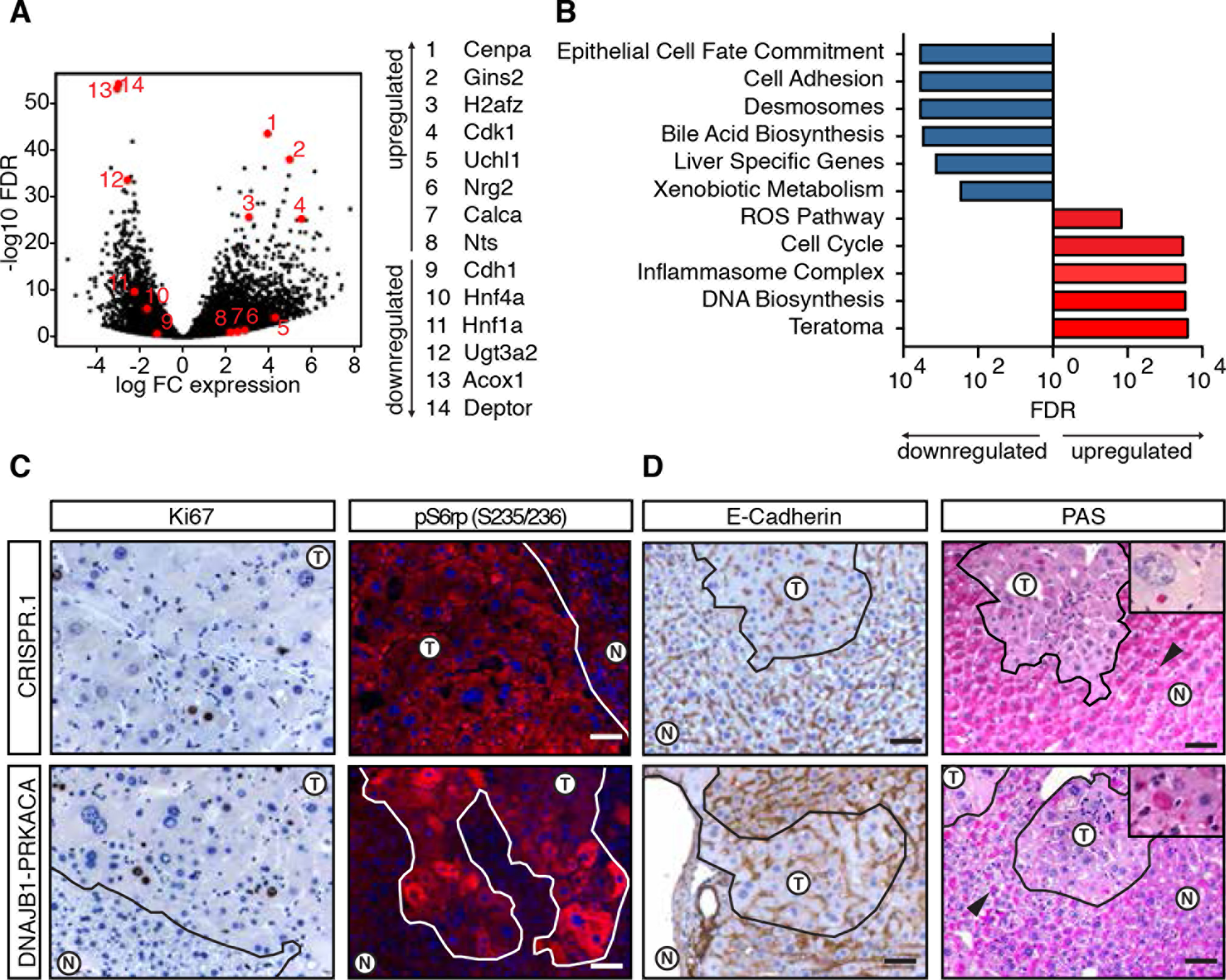
Molecular profiling of murine FL-HCC reveals processes linked to tumorigenesis. (**A**) Volcano plot depicting differentially expressed genes in CRISPR-induced tumors with respect to normal liver. (**B**) ssGSEA analysis for select functionally annotated gene sets. (**C**) Ki67 IHC and p-S6rp S235/236 IF of CRISPR-induced tumor (top), or transposon-induced tumor (bottom). (T: tumor, N: normal) (**D**) E-cadherin (CDH1) staining and Periodic Acid Schiff (PAS) staining. Note adjacent normal hepatocytes positive with asymmetrical subcellular distribution (black arrow). Inset: PAS_+_ Hyaline bodies. All scale bars are 50 αm.

Supporting previous reports that dedifferentiation is associated with *DNAJB1-PRKACA*-driven transformation (32), we observed downregulation of hepatocyte lineage markers and upregulation of some neuroendocrine markers (**Fig. 4D**, **S4G**), as has been observed in human FL-HCC (30). GSEA further showed downregulation of epithelial cell fate commitment, bile acid biosynthesis, liver specific genes, and xenobiotic metabolism (33) and upregulation of a teratoma-associated gene set (**Fig. 4E**). Similarly, gene expression signatures defining specialized zones of hepatocytes (34) were universally lost, consistent with human data (30) (**Fig. S4H**).

Murine FL-HCC cells may also have defects in cell adhesion. Tumor cells displayed a decrease in *Cdh1*/E-cadherin expression and cell adhesion- and desmosome-associated gene sets were downregulated (**Fig. 4A,B**). E-cadherin downregulation was confirmed by IHC and EM revealed indistinct or simple tumor cell-cell junctions (**Fig. S2J,K**). Loss of cell polarity was indicated by a loss of synchronized glycogen storage evident in neighboring hepatocytes, shown by largely negative Periodic Acid Schiff (PAS) staining (**Fig. 4D**). An exception to this pattern was the observation of PAS+ Mallory/hyaline bodies, which are commonly found in FL-HCC (8) and are reminiscent of a stress-induced phenotype (35) (**Fig. 4D**, **inset**).

Murine and human tumors also showed evidence of a response to oxidative stress, as indicated by the upregulation of enzymes involved in detoxifying reactive oxygen species (e.g. *Nqo1*, *Gpx3*, *Gpx4, and Acox1*) (**Fig. 4B**,**S4F**). The accumulation of mitochondria can be driven by oxidative stress (36); upregulation of mitochondrial-encoded transcripts (**Fig. S4B**) and an increase in the number of mitochondria observed by EM (**Fig. 3**,**S2E-H**) were also evident in both the mouse model and clinical samples. Whether or how each of these features contributes to the pathogenesis of FL-HCC remains to be determined.

### WNT pathway cooperates with *DNAJB1-PRKACA* to accelerate FL-HCC

To further understand the genetic basis of FL-HCC and to address the long latency of the single hit models, we investigated additional factors that could accelerate disease. We queried the MSK-IMPACT collection of targeted sequencing data of over 18,000 cancer patients (37). Eighteen liver cancer patients (age 18-36) whose tumors harbored the *DNAJB1-PRKACA* fusion were identified (**Fig. 5A**). As expected, the fusion was not detected in any liver cancer patient over the age of 36 (0/414 patients) or in any non-liver cancer patient (0/18,367 patients), confirming the remarkable specificity of the *DNAJB1-PRKACA* fusion to liver oncogenesis (**Fig. S5A**). While HCC and intrahepatic cholangiocarcinoma (ICC) share several common mutations, none of these have been linked to FL-HCC. Surprisingly, we noted previously unreported recurrent mutations in the Wnt pathway in human FL-HCC (**Fig. 5A,B**). The MSK-IMPACT cohort contains 3/18 (17%) samples of FL-HCC with *CTNNB1* or *APC* mutation (age 18-21 years old). These cases each showed classic histological features of FL-HCC (**Fig. S5C**).

**Fig. 5.**
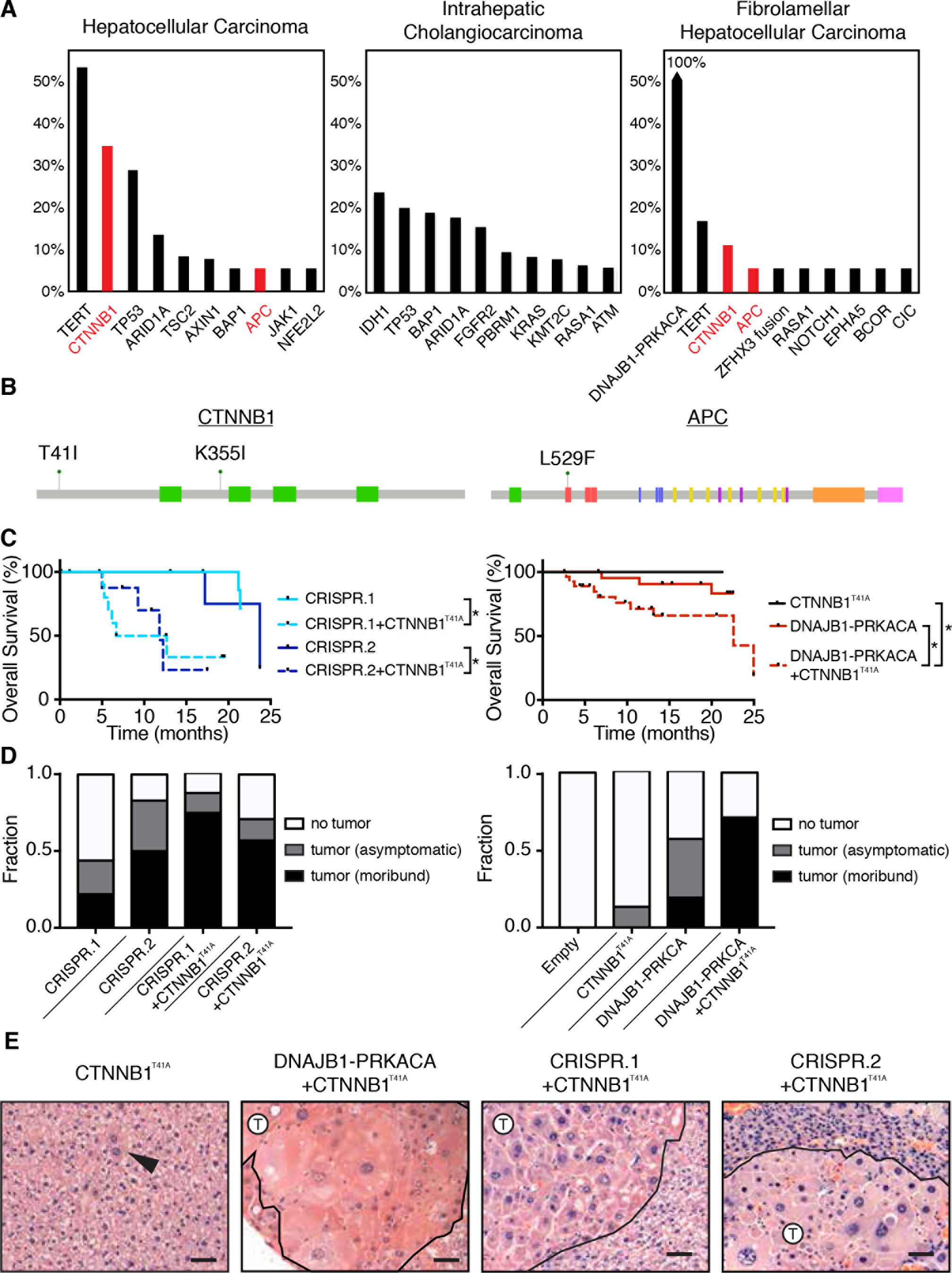
WNT pathway cooperates with *DNAJB1-PRKACA* to accelerate FL-HCC. (A) Frequency of top 10 most commonly mutated genes in hepatocellular carcinoma (HCC, n=142), intrahepatic cholangiocarcinoma (ICC, n=175), and fibrolamellar carcinoma (FL-HCC, n=18) samples from MSKCC IMPACT sequencing. (**B**) Domain schematic of loci of Wnt pathway mutations in FL-HCC. (**C**) Overall survival of mice injected with *CTNNB1^T41A^* and: CRISPR.1 (n=10, log rank p=0.0042 +/− *CTNNB1^T41A^*), CRISPR.2 (n=9, p=0.022 +/− *CTNNB1^T41A^*), and *DNAJB1-PRKACA* (n=30, p=0.069 +/−*CTNNB1^T41A^*). CRISPR.1, CRISPR.2, and *DNAJB1-PRKACA* alone are repeated from (**Figs. 1-2**) for reference. (**D**) Fraction of mice harvested with no detectable tumor, asymptomatic mice with histologically detectable disease (asymptomatic), or moribund mice with tumors (moribund) for each indicated genotype. (**E**) H&E images depicting histology of mice injected with *CTNNB1^T41A^* in combination with CRISPR.1, CRISPR.2, or *DNAJB1-PRKACA*.

In parallel, candidate drivers of liver cancer were evaluated for their ability to synergize with *DNAJB1-PRKACA* to transform hepatocytes. Neither transposon based delivery of *MYC*, *AKT*^myristoylated^, *NOTCH*^ICD^, *YAP*^S127A^, *Fgf15*, *Il10*, *Il18*, nor CRISPR-mediated inactivation or knockout of *p19^ARF^*, *Pten*,
*Prkar1a*, *Rb1*, *Cdkn1b*, and *Tsc2*, cooperated with *DNAJB1-PRKACA* (**Fig. S6**). However, an activated form of β-catenin uniquely cooperated with *DNAJB1-PRKACA* (using a transposon encoding *CTNNB1*^T41A^ cDNA). Of note, the *CTNNB1^T41A^* is the same allele that co-occurred with the *DNAJB1-PRKACA* fusion in a human primary FL-HCC and its corresponding brain metastasis (**Fig. 5B**, **Fig. S5B**). Both pairs of tandem guide CRISPRs, as well as transposon delivery of fusion cDNA, synergized with transposon delivery of stabilized β-catenin, increasing penetrance and reducing latency of the model (**Fig. 5C,D**). In all cases, the histology of the resulting tumors matched the single hit models, though some features (e.g. cell size) were more pronounced (**Fig. 5E**). The acceleration of the model by Wnt signaling was further validated by the combination of *DNAJB1-PRKACA* cDNA and disruption of *Apc* using CRISPR, which yielded tumors with a similar phenotype (**Fig. S7**). While expression of the *DNAJB1-PRKACA* fusion alone led to increased membranous β-catenin, and phosphorylation of β-catenin at PKA phosphorylation site S675, expression of the canonical Wnt target AXIN2 was negative or weak in samples without genetic manipulation of the Wnt pathway (**Fig. S7**), which is corroborated by low expression of AXIN2 mRNA (**Table S2**) and the “HALLMARK_WNT_BETA_CATENIN_SIGNALING” gene set (**Table S3**) in samples driven by the fusion only. Of note, tumor cells arising in a mouse injected with CRISPR.1 sgRNA pair and the *CTNNB1*^T41A^ transposon formed tumors upon multiple rounds of subcutaneous transplantation into syngeneic recipients (**Figure S8**). Hence, genetic lesions that activate Wnt signaling occur in the human disease and can cooperate with *DNAJB1-PRKACA* to accelerate FL-HCC.

### Inflammatory and fibrotic agent DDC enhances FL-HCC tumorigenesis

FL-HCC typically occurs in young patients that do not have the overt chronic liver diseases that often promote fibrosis and contribute to the emergence of classic HCC (8). Nevertheless, since the tumors arising in our mouse model lacked the eponymous fibrosis characteristic of human FL-HCC, we wondered whether experimental strategies to induce fibrosis might accelerate murine tumors. The hepatotoxin 3,5-diethoxycarbonyl-1,4-dihydrocollidine (DDC) causes oxidative liver damage, cell death in periportal hepatocytes, atypical ductal expansion of progenitor cells, and ultimately, fibrosis (38), and can accelerate HCC tumorigenesis by specific oncogenic events (39, 40).

Consistent with published results, mice treated with a 0.1% DDC-containing diet develop hepatomegaly, inflammation, and fibrosis with portal bridging by 8 weeks of treatment, but did not develop tumors (38, 41) (**Fig. 6A-C**,**S9A**). Although DDC diet showed some increase in the onset of tumors following expression of mutant *CTNNB1* alone, the effects on the combination of mutant *CTNNB1* and *DNAJB1-PRKACA* were dramatic: in fact, 6-month survival was decreased from 60-70% with *CTNNB1^T41A^*/*DNAJB1-PRKACA* to 0% observed with the same combination in DDC-treated mice (**Fig. 6A,B**). The histology of the tumor cells themselves remained largely unchanged by DDC treatment, but as expected, the surrounding tissue acquired DDC-associated phenotypes associated with tissue regeneration following injury (**Fig. 6C**,**S9B**,**S10**). Surprisingly, the morbidity of the combination often preceded the establishment of significant fibrosis (**Fig. 6C**,**S9**). Therefore, these data suggest that one or more factors associated with the DDC-induced regenerative response can fuel murine FL-HCC. Furthermore, the combination of our non-germline genetic approaches and a DDC diet produces FL-HCC like tumors at high penetrance and with a short latency.

**Fig. 6.**
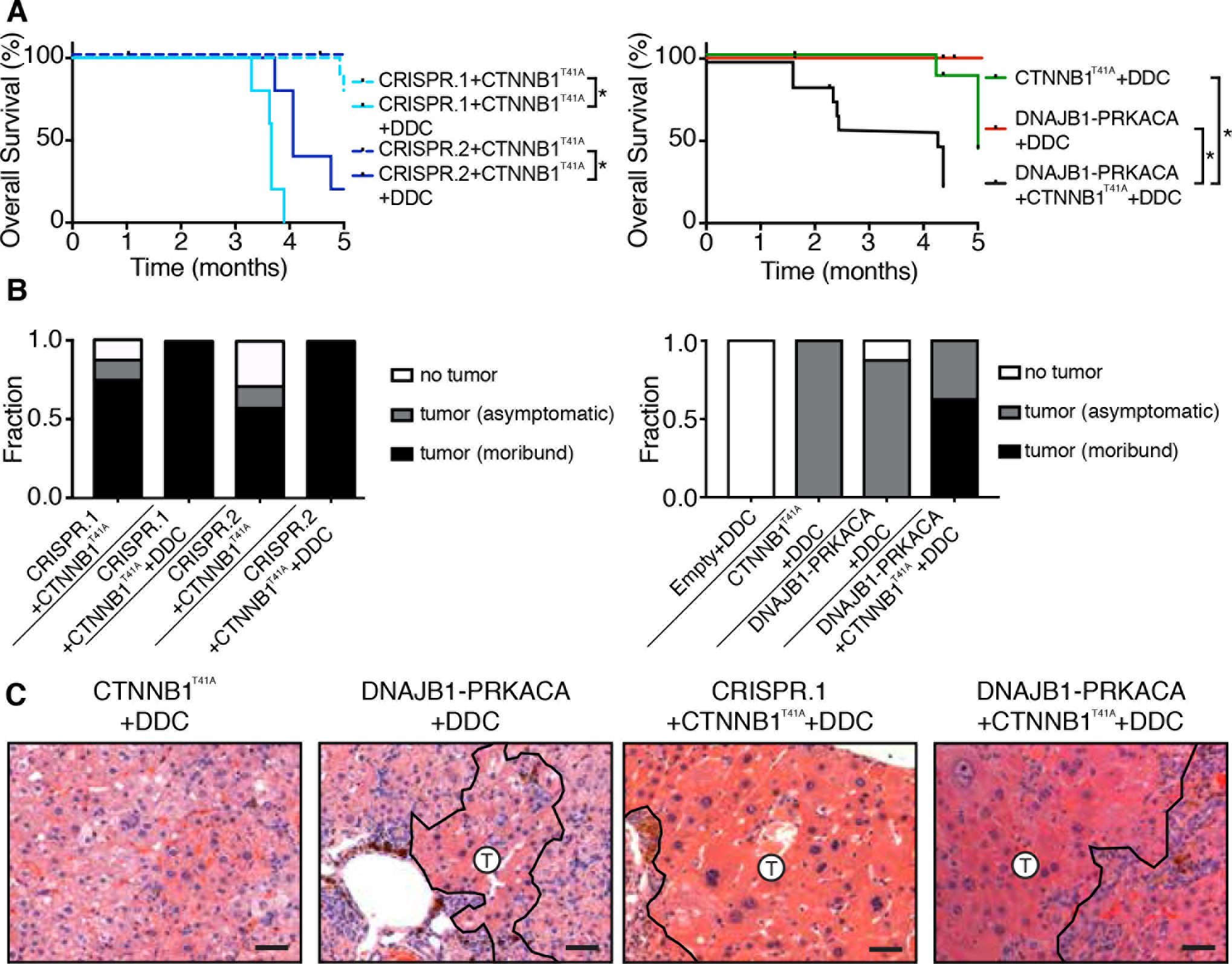
Inflammatory and fibrotic agent DDC enhances FL-HCC tumorigenesis. (**A**) Overall survival of mice injected with indicated genotypes +/- DDC diet: CRISPR.1+ *CTNNB1^T41A^* (n=5, log rank p<0.001 +/- DDC), CRISPR.2+ *CTNNB1^T41A^* (n=5, log rank p<0.001 +/- DDC), *DNAJB1-PRKACA*+ *CTNNB1^T41A^* (n=15, log rank p=0.001 +/- DDC). Survival data of mice without DDC treatment repeated from (**Figs. 1,2,5**). (**B**) Fraction of mice harvested with no detectable tumor, asymptomatic mice with histologically detectable disease (asymptomatic), or moribund mice with tumors (moribund) for each indicated genotype with or without DDC diet. (**C**) H&E images depicting histology of mice of the indicated genotypes and fed DDC diet.

### Tumorigenicity of DNAJB1-PRKACA is dependent on kinase domain

To illustrate the potential of a rapid and robust model of FL-HCC, we used the above methods to address whether the kinase activity of the *DNAJB1-PRKACA* fusion is essential for its ability to drive tumorigenesis—a prerequisite for rationalizing the use of small molecule inhibitors targeting the PRKACA kinase for treatment of FL-HCC. To this end, we produced a kinase dead version of the *DNAJB1-PRKACA* fusion harboring a mutation in the PRKACA component, equivalent to the previously described K72H mutation (42), and compared its oncogenic potential to the intact fusion cDNA when combined with DDC and *CTNNB1*^T41A^. A cohort of mice was produced and sacrificed at 9-10 weeks to examine the presence or absence of liver lesions. While clusters of neoplastic hepatocytes with classic FL-HCC morphology were observed in samples with an intact kinase domain, no such atypical hepatocytes were identified in samples expressing the kinase dead (KD) fusion cDNA (**Fig. 7A**). Furthermore, expression of the kinase intact fusion led to a significantly elevated Ki67 positive fraction of GFP_+_ cells, while the GFP_+_ cells in samples expressing the kinase dead fusion cDNA did not show a significantly higher Ki76 positive fraction compared to adjacent normal liver or DDC-treated empty vector controls (**Fig. 7C**). Thus, the PRKACA kinase domain is required for FL-HCC tumor initiation.

**Fig. 7.**
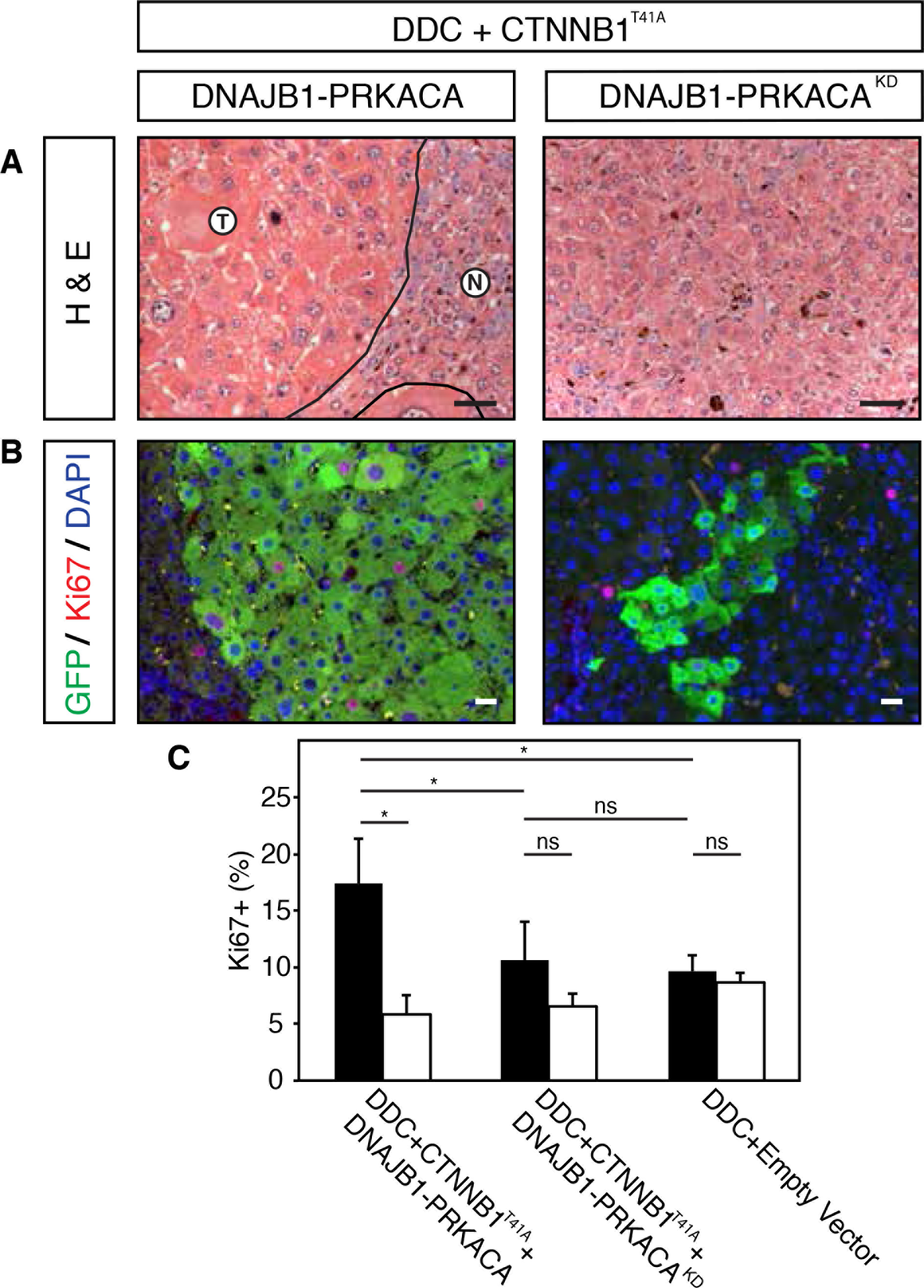
Tumorigenicity of DNAJB1-PRKACA is dependent on kinase domain. (**A**) H&E staining of DDC-treated mouse livers transfected with *CTNNB1^T41A^* combined with either *DNAJB1-PRKACA* or kinase-dead *DNAJB1-PRKACA* (KD, right) (**B**) Immunofluorescence co-staining for Ki67 (red) and GFP (green). (**C**) Quantification of Ki67 positive cells. Error bars represent standard deviation. Two-tailed t-test p<0.05 (**\***).

## Discussion

Using a combination of hydrodynamic transfection, somatic genome editing, and transposon-mediated gene delivery, we demonstrate that the *DNAJB1-PRKACA* fusion is a *bona fide* oncogene that drives FL-HCC. Since tumors were produced by both the endogenous fusion and ectopic expression of the fusion cDNA, it appears that the loss of the intervening 400kb deleted region, encompassing 7 additional genes, is dispensable for tumorigenesis. The gene fusion appears to be functionally distinct from the mere overexpression of wild type *PRKACA*, which is insufficient to drive tumor progression, at least when expressed in adult hepatocytes. On the other hand, the kinase domain of PRKACA is required for the FL-HCC phenotype. Importantly, this indicates the potentially druggable enzymatic activity of the chimera is important for tumorigenesis.

Numerous features of the FL-HCC models described here mimic the human disease. Histologically, our methods generate tumors whose morphology is strikingly reminiscent of human FL-HCC. Dramatic accumulation of mitochondria appears to be a consistent consequence of DNAJB1-PRKACA activity, further linking our model to the human disease. Globally, we observe an expression signature significantly enriched with the genes differentially expressed in human FL-HCC. Nonetheless, DNAJB1-PRKACA expression in the murine hepatocytes does not appear to directly drive all molecular features of FL-HCC (fibrosis, expression of biliary markers, CD68). Additionally, while human FL-HCC often metastasizes, no metastases were observed in the mouse model. Perhaps other biological factors, or time, are needed for metastatic progression. These differences notwithstanding, the mouse model allows for insight into the phenotype of protein kinase A fusion activity in the liver in a controlled setting. We observe that induction of the fusion results in tumors with activated mTOR signaling (a druggable pathway) and detectable effects of oxidative stress (a potential vulnerability), validated by multiple previous reports describing human clinical samples (7, 30, 31, 33).

In human FL-HCC, we observed recurrent mutations that hyperactivate the Wnt pathway together with the DNAJB1-PRKACA fusion. Furthermore, genetic alteration of this pathway – but not several other oncogenes or tumor suppressors – synergized with DNAJB1-PRKACA-driven tumorigenesis in the mouse. While the basis for this genetic interaction remains to be determined, it is possible that modification of β-catenin may be one downstream output of DNAJB1-PRKACA and stabilization of β-catenin may amplify its functional consequences. PKA has been described to regulate β-catenin through a variety of mechanisms, including direct modification via C-terminal phosphorylation (43-45); accordingly, phospho-β-catenin is elevated in human FL-HCC (46, 47). Alternatively, β-catenin may protect cells from oxidative stress imposed by DNAJB1-PRKACA (48).

DDC dramatically shortened tumor latency, unexpectedly preceding extensive fibrosis. By causing oxidative stress-induced cell death in a subset of periportal hepatocytes, DDC sets in motion a liver regenerative response that is associated with compensatory expansion of liver progenitor cells, activation of myofibroblasts, fibrosis, and massive immune cell infiltration (41) in a process supported by β-catenin expression in hepatocytes (48). Any or all of the activated and recruited cell types could contribute paracrine signals that protect and stimulate growth in the spatially separated (largely pericentral) population of hepatocytes that are transfected by hydrodynamic injection (49). While chronic liver damage is not considered to be necessary for human FL-HCC, our data raise the possibility that some environmental factor, perhaps in a susceptible population, may be relevant in the etiology of the disease. Alternatively, it is possible that β-catenin and/or DDC facilitated a change in cell state that supports or expands a susceptible progenitor cell population more prevalent in adolescents.

Animal models set a gold standard for assessing the oncogenic potential of aberrations observed in human cancer and provide experimental systems to study disease mechanisms or test novel therapeutic strategies (14). While one patient-derived xenograft (PDX) of FL-HCC has been described (32), our models introduce the ability to examine the entire process of tumor initiation in immune competent organisms. The somatic engineering methods described here involve only delivery of plasmid DNA to hepatocytes and do not require expensive and time-consuming generation or breeding of germline mouse strains, cell transplantation, or stable expression of Cas9. The model is easily implemented, reproducible, and genetically defined. As such, these systems are a powerful platform to further understand the biology of FL-HCC and facilitate drug discovery for a disease that disproportionately affects young patients and has limited treatment options.

## Materials and Methods

### Vectors and cloning

One sgRNA was cloned into the px330 vector, which was a gift from Feng Zhang (Addgene plasmid #42230), and a second U6-sgRNA cassette was inserted in the XbaI site with XbaI-NheI overhangs. As indicated, for other experiments (**Fig. S1**), the lenti-CRISPR vector was a gift from David Sabatini (Addgene plasmid #70662). The pT3 transposon and SBase vectors were a kind gift of Dr. Xin Chen, University of California at San Francisco. Sequences for sgRNAs, cDNAs and primers are listed in **Supplementary Table 4**.

### Animals and Treatments

Female, 6- to 10- week-old C57BL6/N mice were purchased from Envigo (East Millstone, NJ). All animal experiments were approved by the MSKCC Institutional Animal Care and Use Committee (protocol 11-06-011). For hydrodynamic tail-vein injection, a sterile 0.9% NaCl solution was prepared containing plasmid DNA of either 40μg CRISPR vector or 20μg transposon vector together with CMV-SB13 Transposase (1:5 molar ratio). Mice were injected into the lateral tail vein with a total volume corresponding to 10% of body weight (typically 2 ml for a 20g mouse) in 5–7 seconds (15, 50). DDC treatment was administered through a diet containing 0.1% DDC (Sigma-Aldrich, St. Louis, Missouri; Envigo, Madison, WI) until sacrifice (40). Transplants were performed by finely mincing freshly isolated tumors, suspending in 1:1 PBS:matrigel, and injecting subcutaneously in a 100 μl volume.

### Electron Microscopy

Tissue was fixed in 4% glutaraldehyde and transferred to cold PBS until further processed. The tissues were post fixed in 1% osmium tetroxide in PBS. After washing in water, the tissue was stained with 2% aqueous uranyl acetate for ~2 h at 4°. Tissues were dehydrated through a series of acetones and propylene oxide and embedded in Epon. Ultrathin sections were deposited on grids and stained with uranyl acetate for 15 minutes and lead citrate for 5 minutes.

### Immunohistochemistry and Immunofluorescence

Tissue was prepared for histology by fixing in 10% buffered formalin overnight then transferred to 70% ethanol until paraffin embedding and sectioning (IDEXX RADIL, Columbia, MO). Antigen retrieval was performed in a pressure cooker with Sodium Citrate buffer. The following primary antibodies were used: Ki67 (Abcam ab16667, 1:200), p-S6rp (Cell Signaling 2211, 1:200), E-cadherin (BD 610181, 1:500), HNF1a (Santa Cruz sc-6547, 1:100), HNF4a (Abcam ab41898, 1:200), CK7 (Abcam ab181598, 1:500), CK19 (Abcam ab133496, 1:1000), CD68 (MSKCC Pathology Core Facility), IBA1 (Wako, 1:500), GFP (Abcam ab13970, 1:200), β-catenin (BD610154, 1:500), p-β-catenin (Cell Signaling 9567,1:100), and AXIN2 (Abcam ab32197, 1:800). Primary antibodies were incubated at 4°C overnight in blocking buffer. Sections were incubated with anti-rabbit ImmPRESS HRP-conjugated secondary antibodies (Vector Laboratories, #MP7401) and chromagen development performed using ImmPact DAB (Vector Laboratories, #SK4105). Stained slides were counterstained with Harris’ hematoxylin. Images of stained sections were acquired on a Zeiss Axioscope Imager Z.1. Raw .tif files were processed using Photoshop CS5 software (Adobe Systems Inc., San Jose, CA) to adjust white balance.

### RNAseq

Total RNA was isolated from frozen tissue using the RNeasy Mini Kit (Qiagen), quality control was performed on an Agilent BioAnalyzer, 500 ng of total RNA (RNA integrity number > 8) underwent polyA selection and Truseq library preparation according to instructions provided by Illumina (TruSeq RNA Sample Prep Kit v.2) with 6 cycles of PCR. Single-end, 75-bp sequencing was performed at the CSHL core facility. Approximately 8 million reads were acquired per sample. Resulting RNA-Seq data was analyzed as described previously (23). Adaptor sequences were removed using Trimmomatic. RNA-seq reads were then aligned to the mouse genome (mm10) using STAR (51) with default parameters and genome-wide transcript counting was performed using subread to generate a count matrix (52, 53). Differential expression analysis was performed by DESeq2 (54). Genes were considered to be significantly differentially expressed if tumor/normal comparison was greater than 2-fold and FDR-adjusted p value was less than 0.05.

Human-Murine mapping of orthologs was performed based on the Ensembl database accessed through the Biomart R/Bioconductor package (55). Human fibrolamellar HCC signatures were defined as genes with at least 2-fold and significant expression changes in fibrolamellar tumors with respect to normal. Human transcriptional profiling data was obtained from published studies (7, 30), and the TCGA/Broad GDAC firehose using annotation from Dinh et al (10). For comparison to human datasets and for gene set enrichment analysis, the ssGSEA method was implemented using the GSVA package within R (29). The GSVA outputs were subsequently compared across groups using the limma package (56). The C2, C3, C5, and Hallmark collections of gene sets from MSigDBv6.0 were queried (57).

### Human Tumor Sequencing Data

The MSK-IMPACT sequencing data (37) were obtained from the MSKCC cBioPortal (58) (http://www.cbioportal.org). Of the 18 *DNAJB1-PRKACA* fusion cases, 14 were annotated as fibrolamellar HCC and 4 were annotated as HCC. We considered all *DNAJB1-PRKACA* positive liver cancers as FL-HCC, given the common of misdiagnosis of this rare cancer type (10), for which the presence of the *DNAJB1-PRKACA* should be considered diagnostic (2).

## Supplementary Figure Legends

**Fig. S1.**
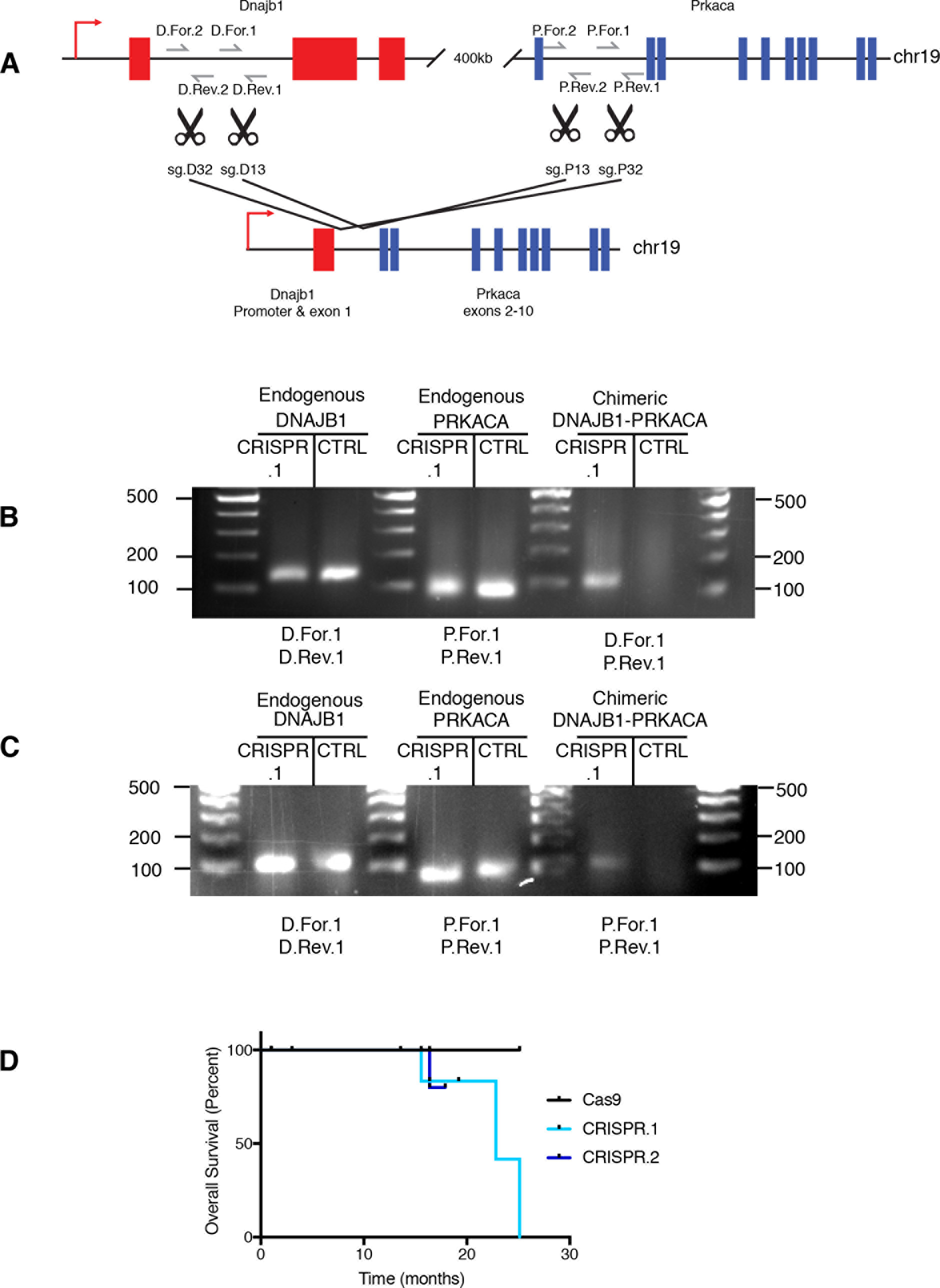
Validation of CRISPR-mediated deletion. (**A**) Schematic of sgRNAs and primers. (**B**) Detection of genomic deletion in NIH-3T3 cells infected with tandem guide lentiCRISPR construct. (**C**) Detection of deletion in genomic DNA extracts from whole livers 4 days following hydrodynamic tail vein injection of the same construct. (**D**) Overall survival of mice following hydrodynamic tail vein injection of the lentiCRISPR plasmid DNA.

**Fig. S2.**
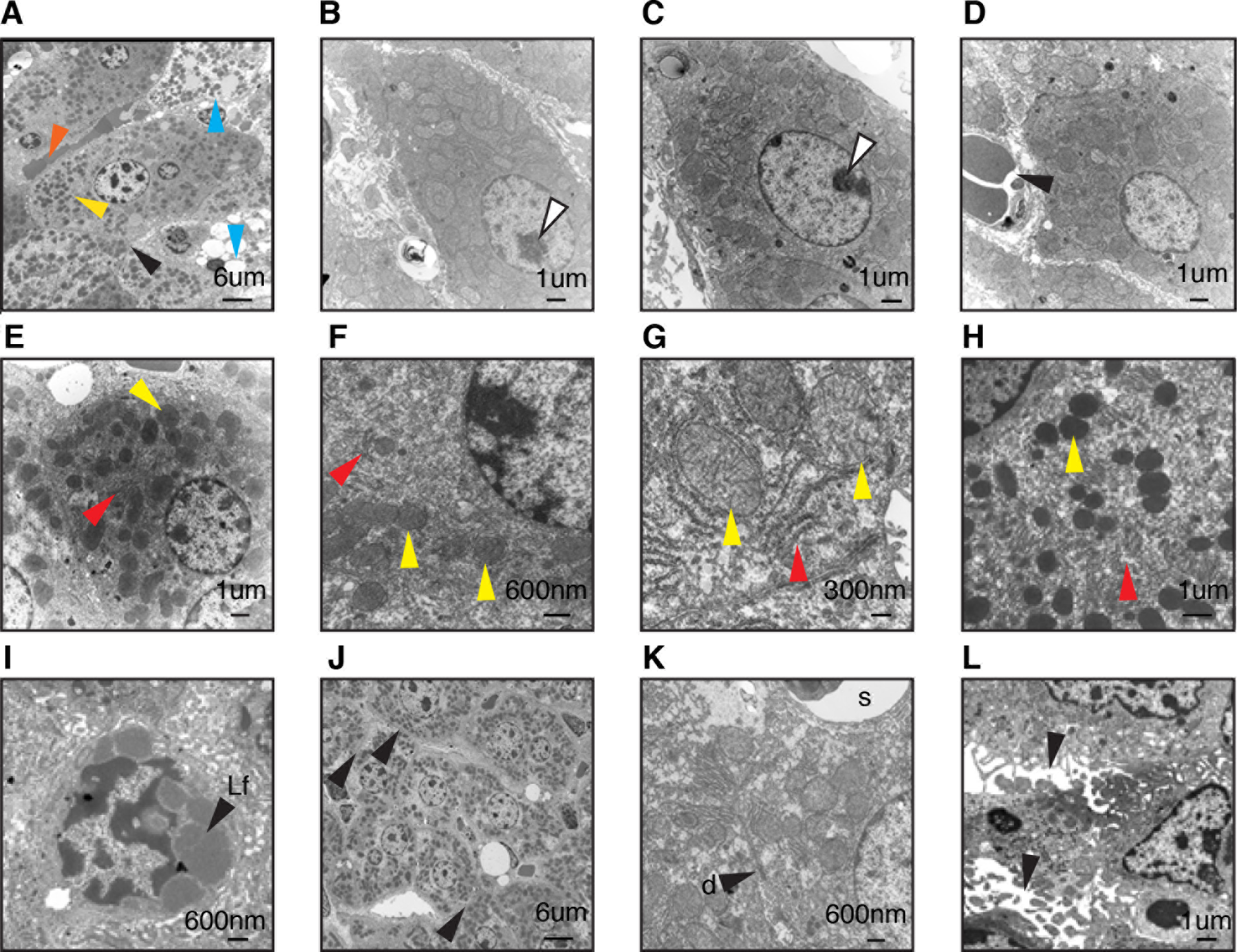
Additional ultrastructural analysis of murine FL-HCC. (**A**) FL-HCC tumor cells are enlarged with abnormally abundant mitochondria (yellow arrowhead). Necrotic cells (blue) and nonfenestrated vessels (orange), and indistinct cell-cell junctions (black). (**B,C**) Tumor nuclei are large and round to indented with prominent nucleoli (white arrows) (**D**) tumor-associated vasculature (black arrow) (**E-H**) Abundant mitochondria (yellow arrows) surrounded by rough endoplasmic reticulum (red arrows) with scant smooth endoplasmic reticulum (**I**) Lipofuscin “Lf” (**J**) Simple, indistinct cell-cell junctions (black arrows) (**K**) rare desmosome “d” with nearby sinusoid “s” (**L**) Bile canniculi are sometimes widened (black arrows).

**Fig. S3.**
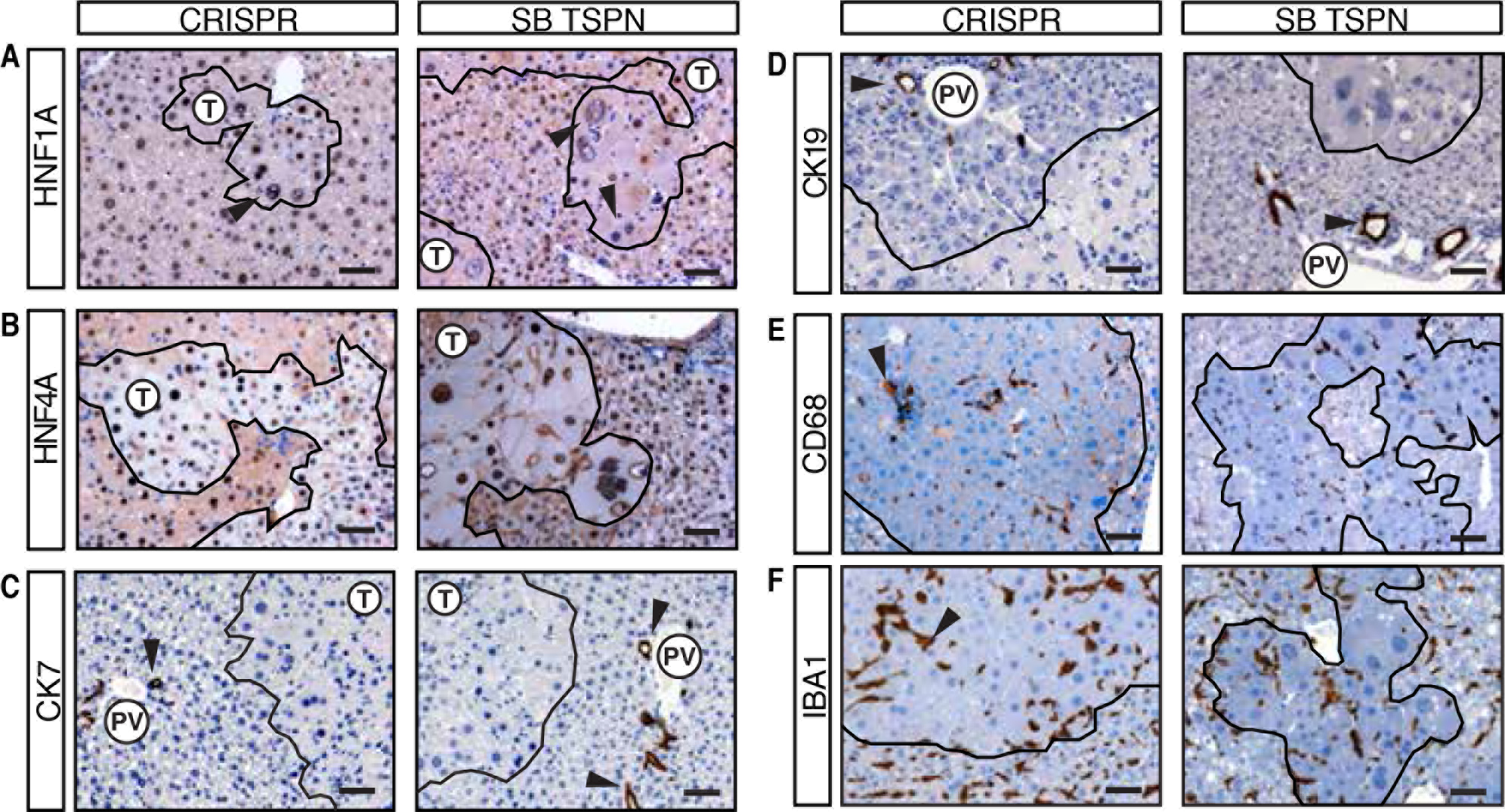
Additional molecular characterization of murine FL-HCC. Indicated genotypes stained for hepatocyte markers (**A**) HNF1A and (**B**) HNF4A, cholangiocyte markers (**C**) CK7 and (**D**) CK19 (arrows: bile ducts, PV:portal vein), (**E**) CD68 (arrows: CD68+ infiltrating macrophages) and (**F**) IBA1 confirming **S3E**. All scale bars are 50 μm.

**Fig. S4.**
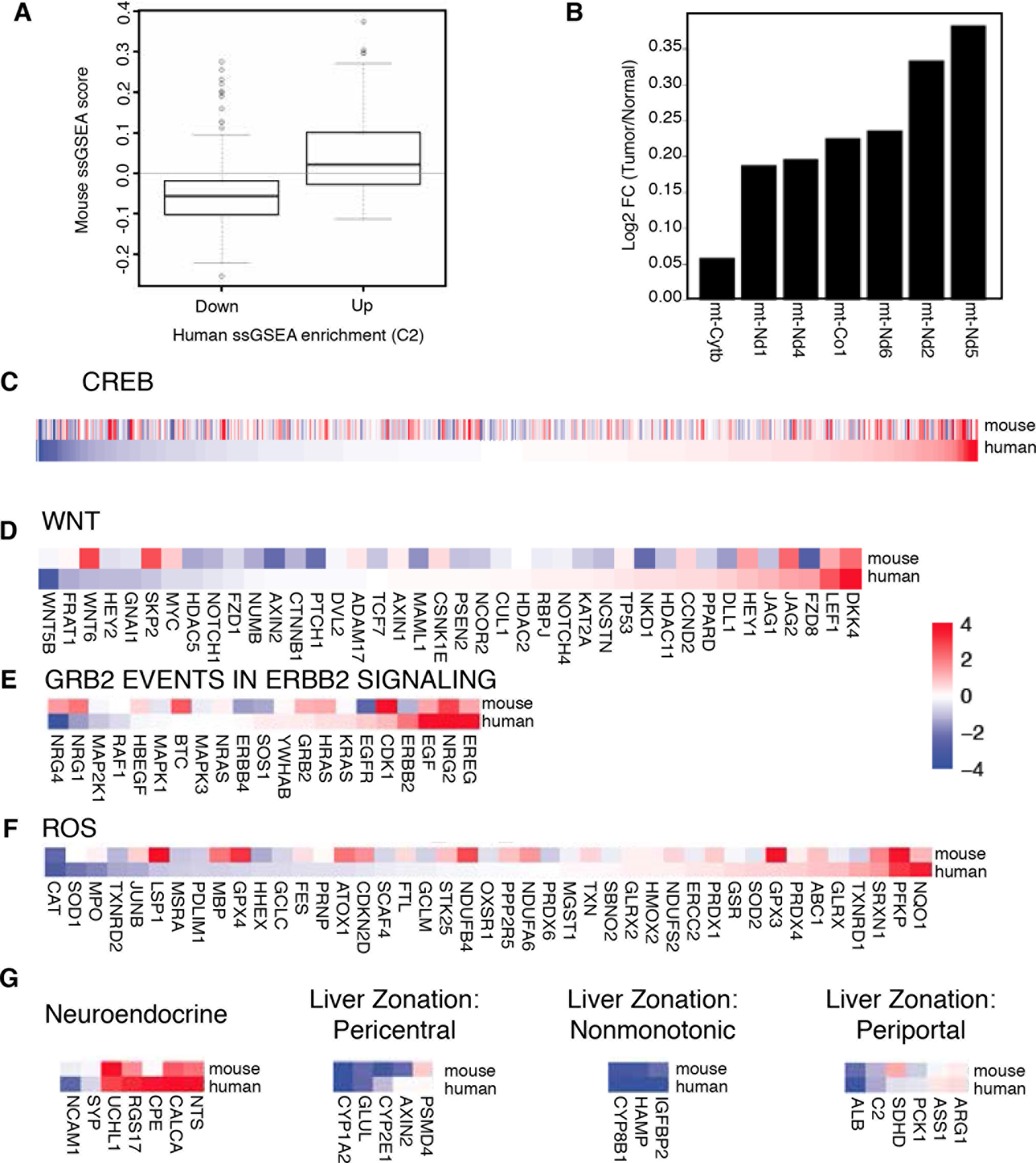
Gene expression of FL-HCC-associated gene sets of interest. (A) ssGSEA enrichment scores from mouse tumor differential expression for gene sets significantly enriched in either upregulated or downregulated human gene expression data (30) (p=4.22-17). (**B**) Gene expression for genes encoded by mitochondrial DNA. Gene expression of corresponding mouse and human (30) genes in the specified gene sets: (**C**) CREB targets (**D**) Wnt signaling pathway (**E**) GRB2 events in ERBB2 signaling (**F**) genes upregulated by ROS (**G**) Lineage specific genes including neuroendocrine markers (30) and (**H**) liver zone-specific genes (34).

**Fig. S5.**
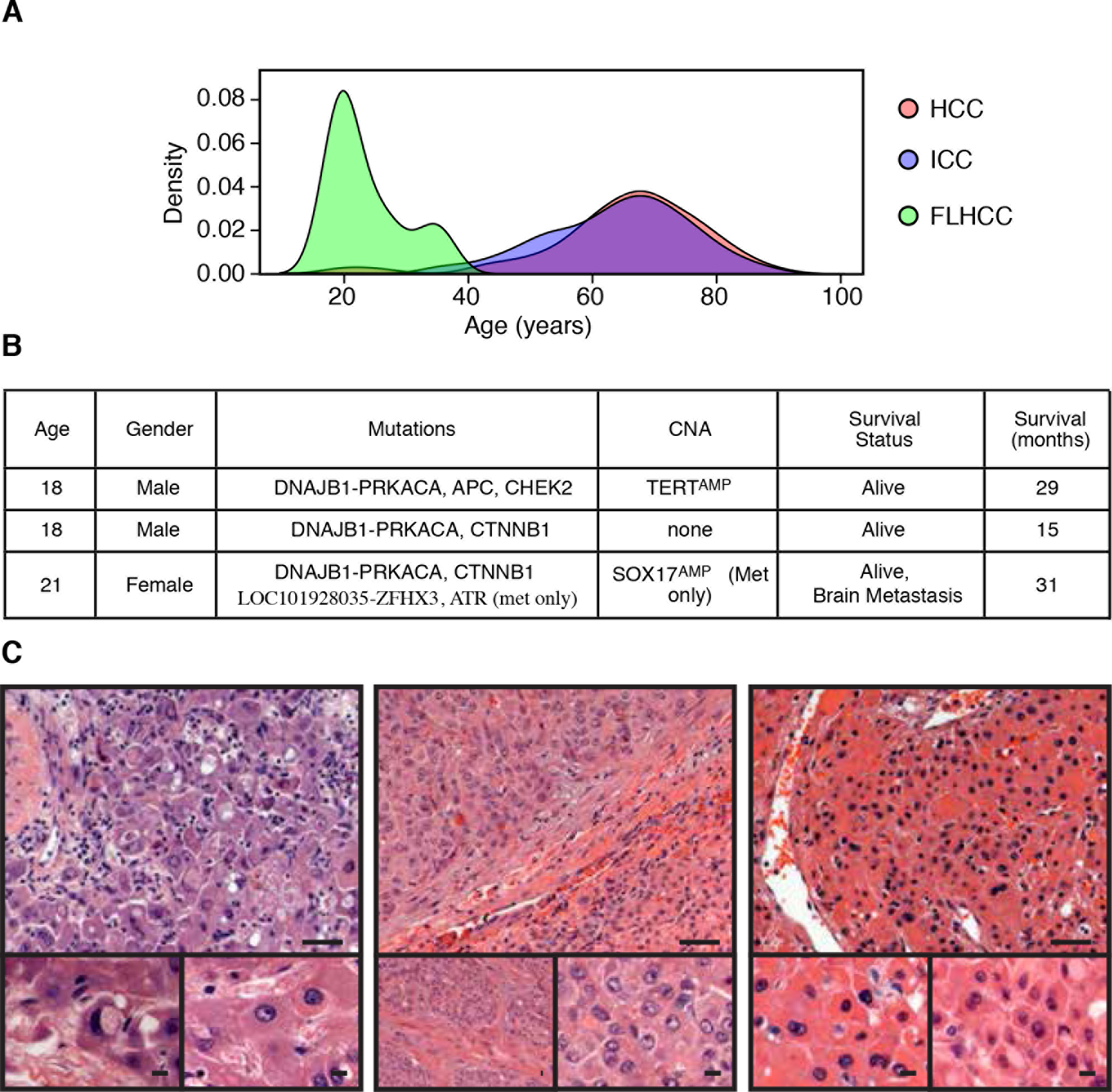
Wnt pathway alteration in human FL-HCC. (**A**) Age distribution of patients with the indicated types of liver tumors (HCC n=129, ICC n=165, FL-HCC n=16). (**B**) Clinical and genomic characteristics of 3 cases of FL-HCC with Wnt pathway mutation. (**C**) H&E staining reveals classic FL-HCC morphology in cases with Wnt pathway mutations. Top: scale bars are 50 μm, bottom (insets): scale bars are 10 μm.

**Fig. S6.**
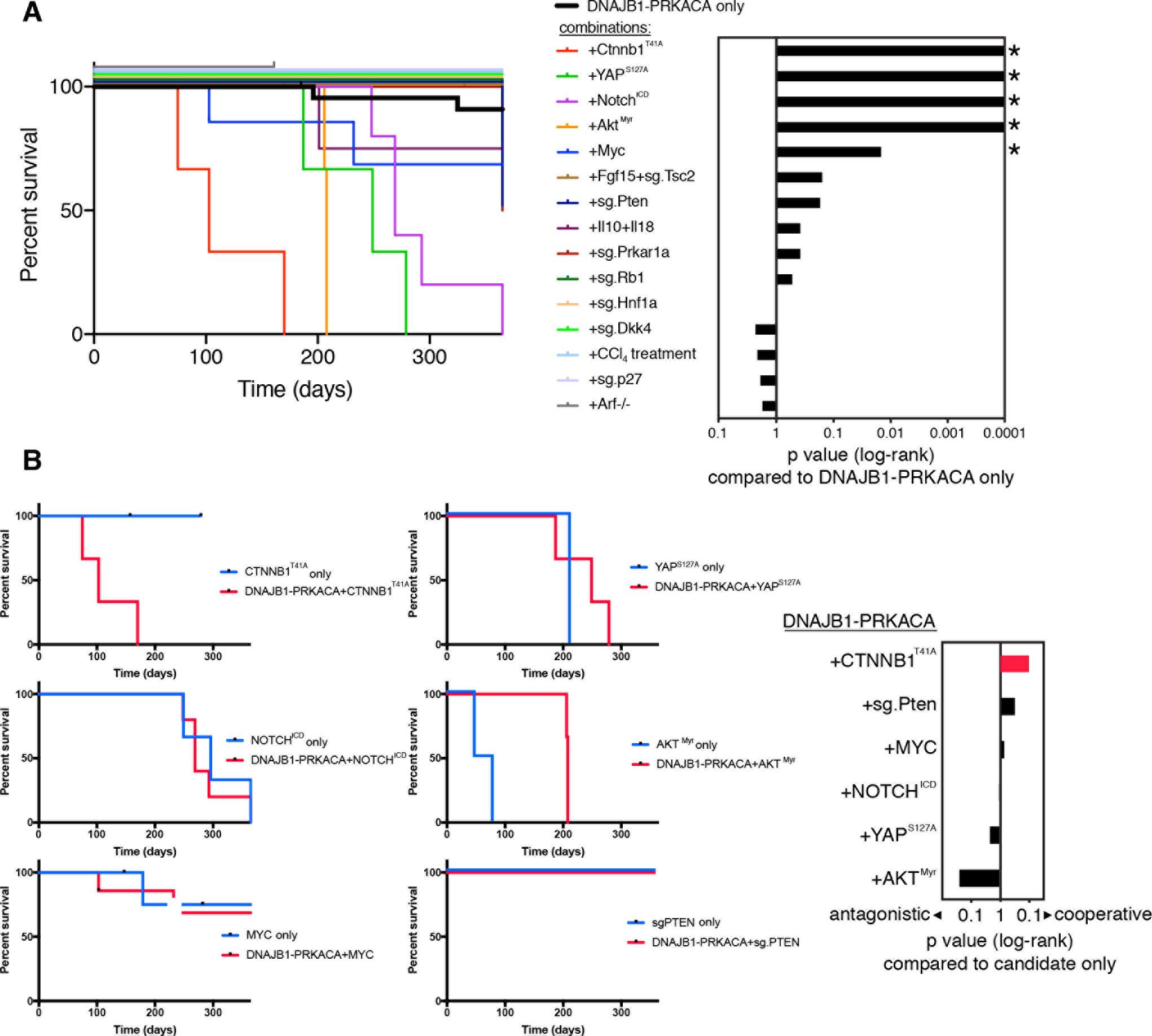
Screen for cooperating mutations. (**A**) Survival data of cohorts injected with DNAJB1-PRKACA fusion cDNA alone or in combination with the indicated genotypes. (**B**) Survival data of cohorts injected with candidate genes from Fig. S6A alone or in combination with the fusion. CTNNB1^T41A^ cooperates with DNAJB1-PRKACA resulting in higher lethality than either gene alone.

**Fig. S7.**
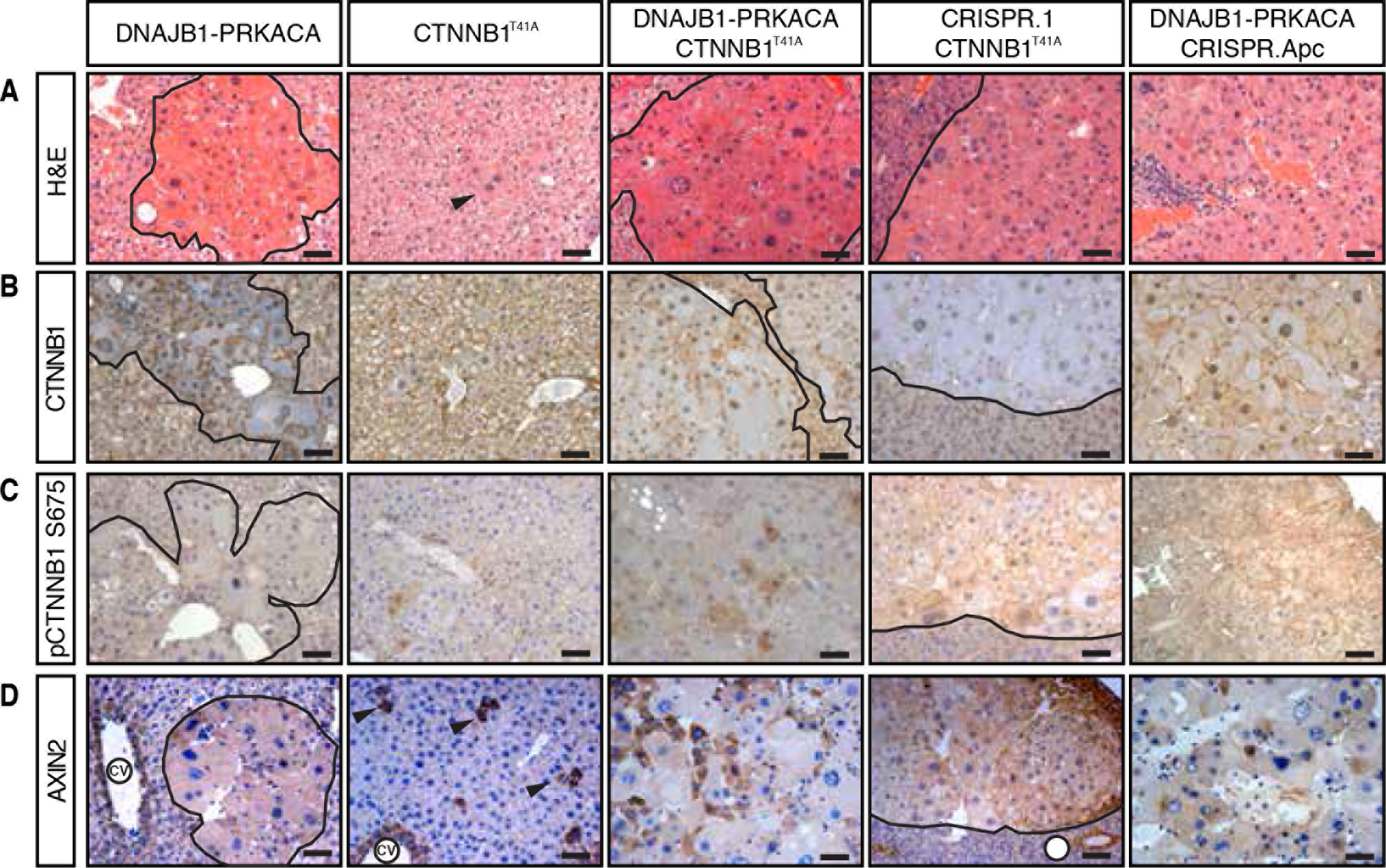
Wnt pathway in murine FL-HCC. Livers injected with *DNAJB1-PRKACA*, *CTNNB1^T41A^*, *DNAJB1-PRKACA + CTNNB1^T41A^*, CRISPR.1 + *CTNNB1^T41A^*, or *DNAJB1-PRKACA +*CRISPR.Apc. (**A**) H&E or immunohistochemistry with antibodies against (**B**) CTNNB1, (**C**) p-CTNNB1 S675 (PKA target site), and (**D**) AXIN2 (Wnt target gene). CV=Central vein (AXIN2 internal positive control). All scale bars are 50 μm.

**Fig. S8.**
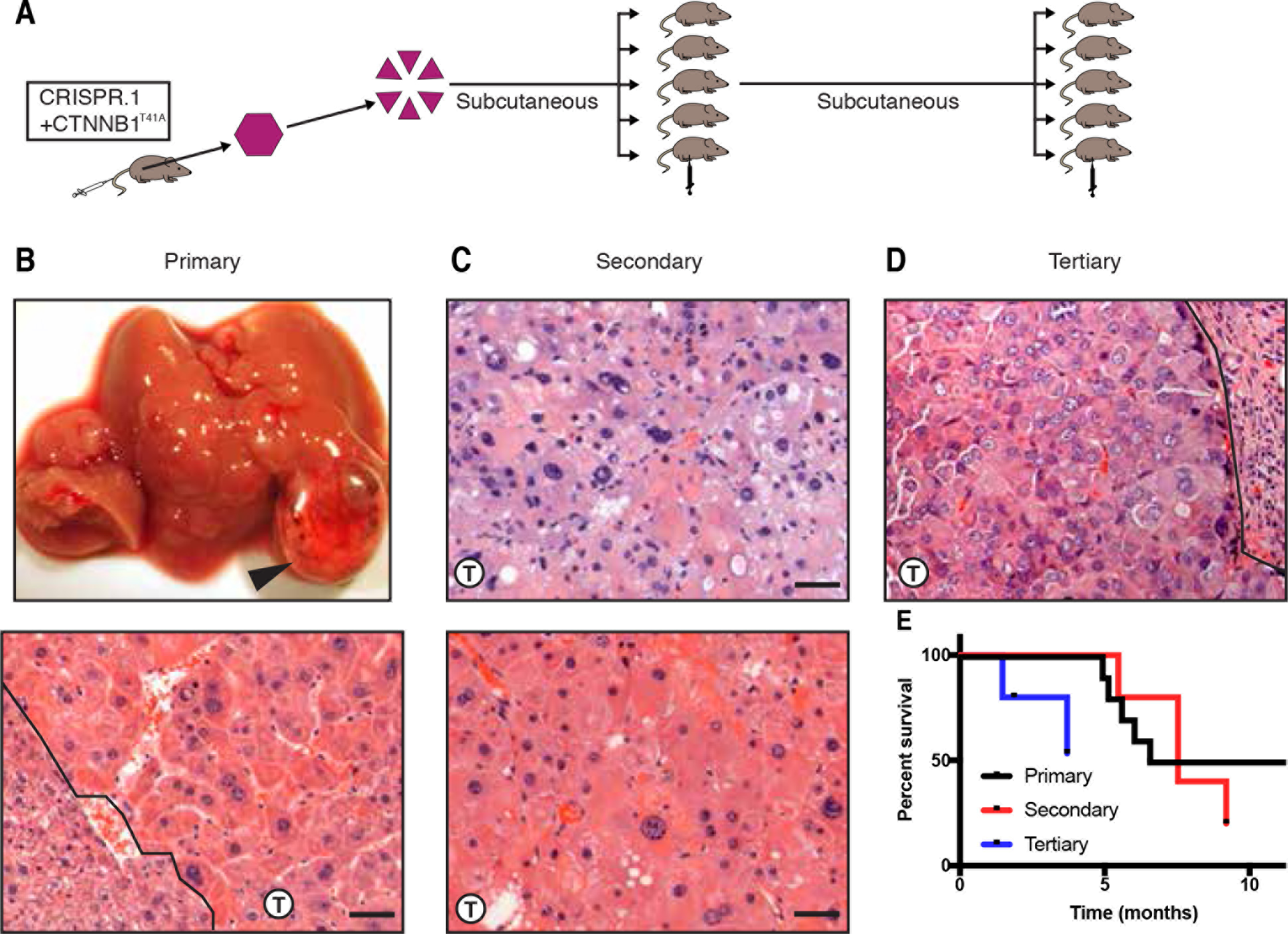
Serial transplantation of murine FL-HCC. (**A**) Primary tumor driven by CRISPR.1 fusion and CTNNB1^T41A^ was minced and transplanted subcutaneously into syngeneic C57/BL6 mice. (**B**) Macroscopic view (top) and histology (bottom) of primary tumor, (**C**) transplanted tumors (first generation) (**D**) and second serial transplantation of a tumor from Fig. S7C (tertiary) (**E**) Survival of cohorts of primary tumors (from Fig. 5C) and tumors from two rounds of serial transplantation.

**Fig. S9.**
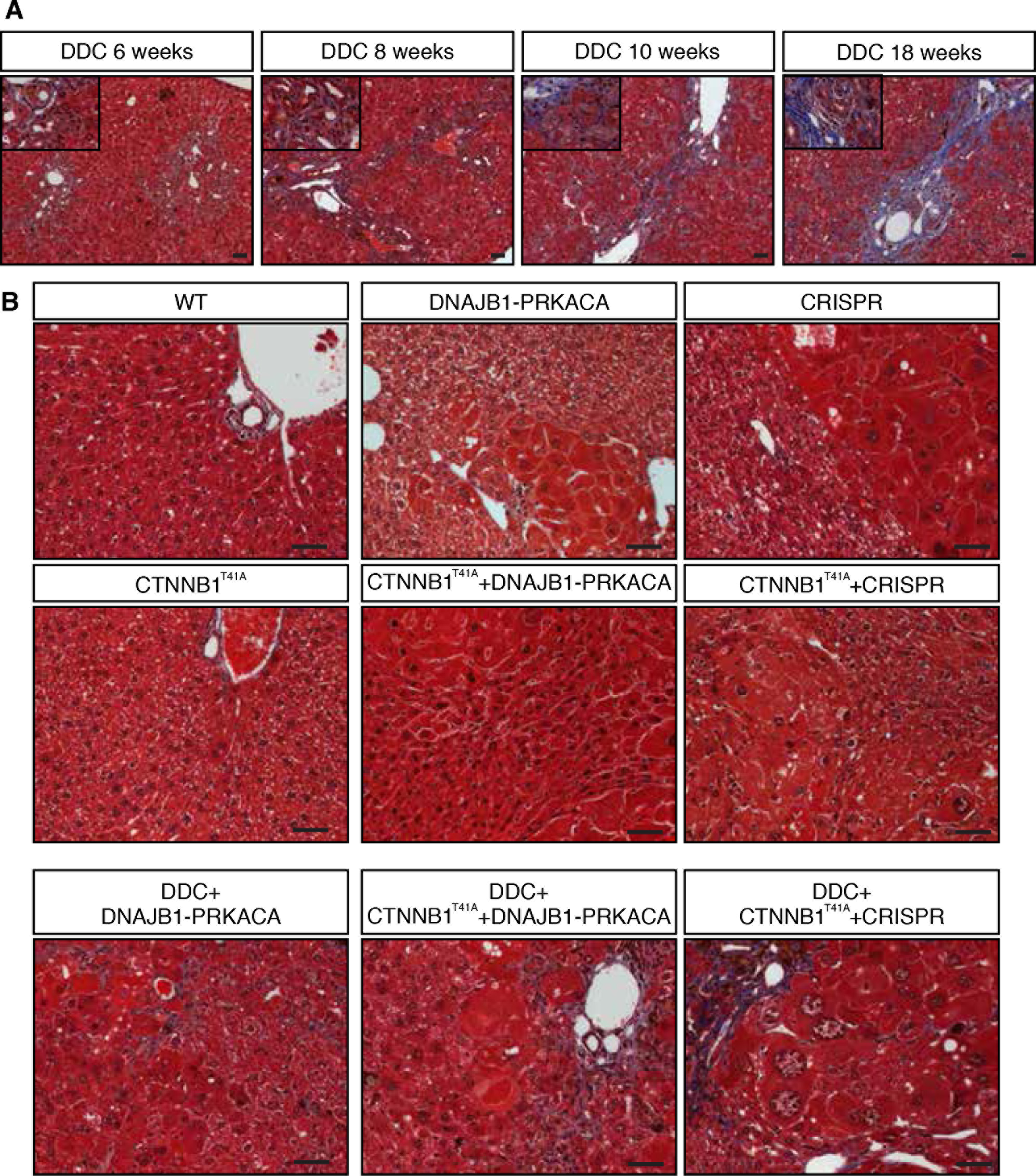
DDC-induced changes in liver pathology. Masson’s Trichrome staining (**A**) indicating progressive development of fibrosis (scale bars: 50 μm) and atypical ductal proliferation (inset). (**B**) No indication of fibrosis was detected in mice with any combination of genetic perturbations, when the mice were fed a normal diet. Mild fibrosis stained positive (blue) in some areas adjacent to tumors following DDC treatment and injection with the indicated constructs.

**Fig. S10.**
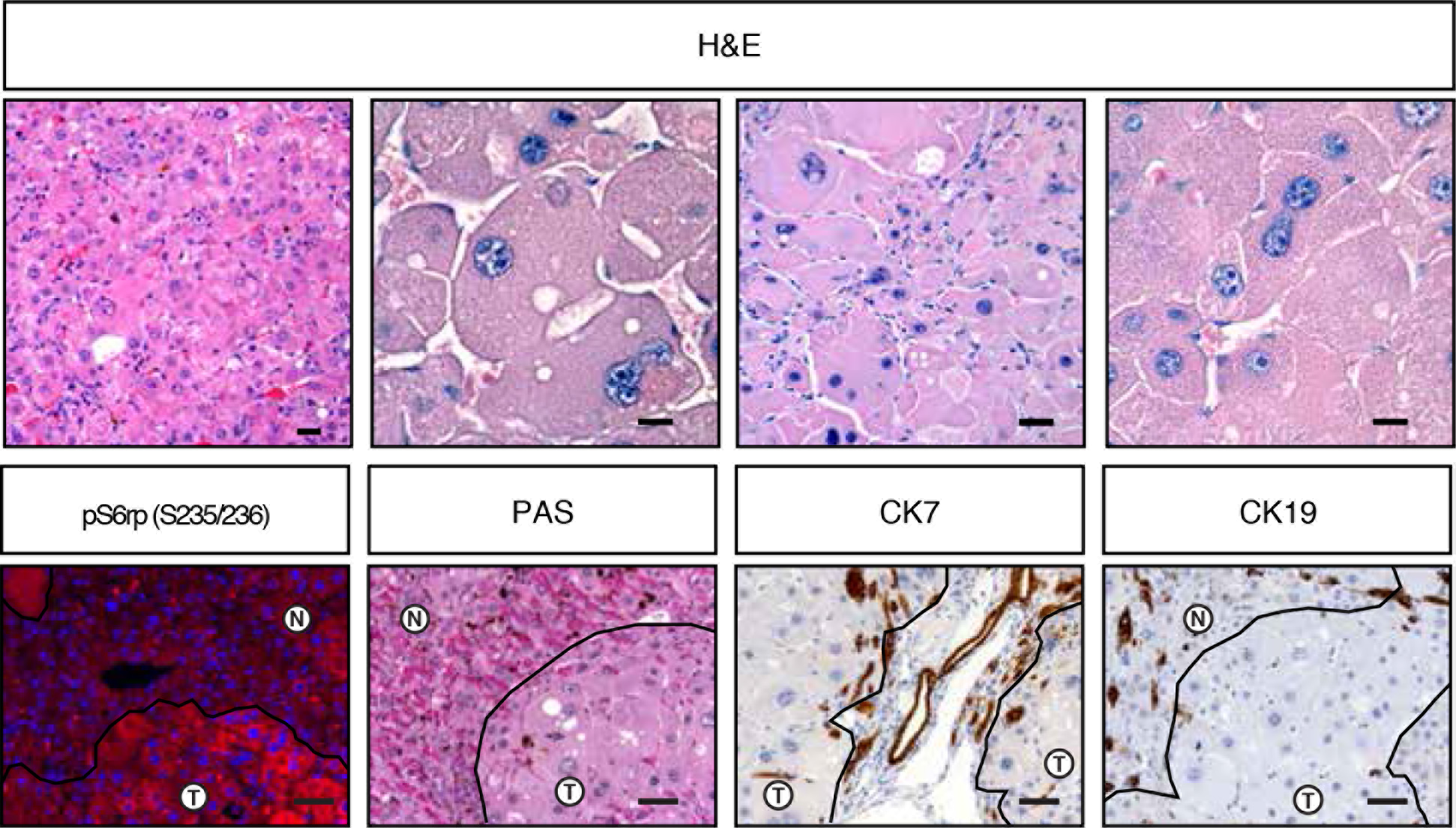
Stable phenotype in DDC-treated tumors. The indicated histological features were found in tumors expressing *DNAJB1-PRKACA* and *CTNNB1*^T41A^ in DDC-treated mice (top). Tumors had recognizable inflammation. Pale bodies, atypical nuclei, and abundant rough endoplasmic reticulum were present. *DNAJB1-PRKACA*/*CTNNB1*^T41A^/DDC tumors were pS6rp^S235/236^+,PAS-, CK17-, and CK19-(bottom).

## Author contributions

E.R.K. performed experiments, analyzed results, performed data analysis and wrote the paper. G.L. and T.B. coordinated experiments and analyzed results. D.F.T., S.L.H., and B.B. provided critical reagents, technical advice and planned experiments. S.T. performed experiments. T.B. performed experiments and performed data analysis of copy number profiling data. C.C. and D.R.E. performed data analysis. J.E.W. performed electron microscopy and provided pathology expertise. S.M.S. conceived the project, supervised experiments, analyzed results, and wrote the paper. S.W.L. conceived the project, supervised experiments, analyzed results, and wrote the paper.

## Acknowledgements

We would like to thank Francisco Sanchez-Rivera (MSKCC), Yadira Soto-Feliciano (Rockefeller University), and John P. Morris IV (MSKCC) for critical reading of the manuscript, and all members of the Lowe laboratory for advice and discussions. We thank Wei Luan and Leah Zamachek for technical assistance and David Klimstra, Paul Romesser, Robert Bowman, Joana DeCampos Vidigal and Andrea Ventura for critical insights and materials. This work was supported primarily by a project grant from the Starr Cancer Consortium (I10-0098) to S.W.L. and S.M.S. This work was funded in part by the Marie-Josée and Henry R. Kravis Center for Molecular Oncology and the National Cancer Institute Cancer Center Core Grant No. P30-CA008748. We gratefully acknowledge the members of the Molecular Diagnostics Service in the Department of Pathology. E.R.K. is supported by an F31 NRSA predoctoral fellowship from the NCI/National Institutes of Health under award number F31CA192835. G.L., D.R. and S.M.S. were supported by a grant from the NIH 1R56CA207929. T.B. is the William C. and Joyce O’Neil Charitable Trust Fellow and is supported by the MSKCC Single Cell Sequenicing Initiative. S.W.L. is an investigator of the Howard Hughes Medical Institute and the Geoffrey Beene Chair for Cancer Biology.

